# Spatial correspondences of Audiovisual Stimuli on Double Flash Illusion Perception and its Cognitive Modeling

**DOI:** 10.64898/2026.02.19.706740

**Authors:** Yabo Zheng, Lihan Chen

## Abstract

Perceptual processing integrates information from multiple sensory modalities to form a coherent representation of the environment. A classic example of such is the Sound-Induced Flash Illusion (SIFI), where the perceived number of visual flashes is altered by conflicting auditory stimuli. While the SIFI is a well-established phenomenon of multisensory integration, the influence of physical spatial characteristics—specifically stimulus eccentricity and spatial congruence—on integration levels remains debated.To address this gap, this study used the SIFI paradigm to investigate the effect of visual stimulus spatial location and the spatial congruence between auditory and visual stimuli on audiovisual integration. In Experiments 1 and 2, we found that when spatial attention was controlled via cueing, unimodal visual performance remained consistent across locations. However, the susceptibility to SIFI increased progressively from the central to the peripheral visual field, exhibiting a spatial pattern of Gaussian distribution. Bayesian modeling further supported this by showing that this spatial modulation was driven by an increase in the integration weight assigned audiovisual representations in the periphery, rather than changes in sensory uncertainty alone. Conversely, Experiment 3 demonstrated that the spatial congruence of audiovisual stimuli did not affect the SIFI or alter the integration processing. These findings refine our current understanding of the spatial modulation upon audiovisual integration. By incorporating the visual system’s spatial properties into a Bayesian framework, we provide a computational explanation for the eccentricity-dependent nature of multisensory integration.

## 1. Introduction

The environment in which we live is rich with information spanning multiple sensory modalities. To facilitate efficient interaction with this environment, the brain adaptively perceives its surroundings by integrating multisensory information (Bauer et al., 2020). Multisensory integration (MSI) is the process by which an observer combines information originating from different sensory channels into a coherent and unified perceptual experience (Stein & Stanford, 2008). This cross-modal integration enhances an observer’s perceptual efficiency and precision, leading to benefits such as reduced reaction times (Pomper et al., 2014), heightened stimulus salience (Driver & Noesselt, 2008), and improved information decoding (Zion Golumbic et al., 2013).

Multisensory integration (MSI) is significantly modulated by visual eccentricity, operating through a complex interplay of spatial and temporal rules where behavioral enhancement, such as faster reaction times and increased detection accuracy, is most robust when stimuli are spatiotemporally congruent (Bruns et al., 2024; Cuppini et al., 2025; Garcia et al., 2017; Porada et al., 2026; Recanzone, 2009). When informational inputs from different sensory modalities conflict, the brain may erroneously integrate mismatched stimuli into a unified percept, resulting in cross-sensory perceptual interference (Sterzer et al., 2009; Wang et al., 2013).As visual eccentricity increases, the auditory localization bias—known as the ventriloquist effect—progressively decreases, a phenomenon consistently observed in both neurocomputational models and empirical data (Cuppini et al., 2025). Other phenomena, such as bistable perception, serve as quantifiable behavioral indicators of an individual’s multisensory processing capacity and their tendency to integrate information (Hirst et al., 2020). A paradigmatic example of this domain is the Sound-Induced Flash Illusion (SIFI), where the high temporal resolution of the auditory channel distorts visual perception (Shams et al., 2002). This illusion typically manifests as “fission”, where a single flash paired with multiple beeps is perceived as multiple flashes (Keil, 2020; Keil & Senkowski, 2018), or “fusion”, where multiple flashes paired with fewer beeps are perceived as a single flash (McGovern et al., 2014). The susceptibility to these illusions is governed by the principle of temporal proximity; stimuli are generally integrated only when they fall within a specific “temporal window of integration” (TWI) (Hirst et al., 2020; Lewald & Guski, 2003; Stein et al., 2014; Stein & Meredith, 1993). Furthermore, this integration process is highly plastic, modulated by factors such as aging (DeLoss et al., 2013), increased cognitive load (Michail & Keil, 2018) and top-down manipulations of perceptual expectations (Wang et al., 2019), all of which can significantly widen or narrow the temporal scale of multisensory integration.

Theoretical frameworks for multisensory integration have evolved from the directed attention hypothesis, which emphasizes attentional resource allocation (Welch et al., 1986) but struggles to explain the automatic nature of cross-modal influences (Odegaard et al., 2016), to the modality appropriateness theory, which posits that sensory dominance is determined by a modality’s precision for a given task (Andersen et al., 2004; Hirst et al., 2020; McGovern et al., 2016).

These cognitive models are complemented by computational approaches, such as maximum likelihood estimation and Bayesian causal inference, which suggest the brain performs optimal perceptual inferences by weighting sensory reliability and assessing common signal sources (Ernst & Bulthoff, 2004; Shams & Beierholm, 2010). Physiologically, these processes are supported by neural oscillation synchronization, where cross-modal communication occurs through phase reset or neural entrainment (Bauer et al., 2021; Fries, 2015; Lakatos et al., 2019; Senkowski & Engel, 2024; Thorne & Debener, 2014). Despite these advancements, significant debate remains regarding the role of spatial characteristics in the Sound-Induced Flash Illusion (SIFI). While neuroimaging suggests enhanced auditory-visual connectivity in the peripheral visual field (Eckert et al., 2008; Ghazanfar & Schroeder, 2006; Rockland & Ojima, 2003), behavioral evidence for this “eccentricity effect” is inconsistent: several studies report increased SIFI susceptibility in the periphery (Chen et al., 2017; Shams et al., 2002; Tremblay et al., 2007), yet others find no such spatial influence (Gavin et al., 2022), particularly at eccentricities yet to be fully explored in humans (Falchier et al., 2002). Furthermore, the interaction between spatial proximity (Stein et al., 2014) and inverse effectiveness (Holmes, 2009) remains unresolved, as empirical results vary on whether spatial disparity modulates or has no effect on illusion perception (Aller et al., 2015; Innes-Brown & Crewther, 2009).

Therefore, the impact of stimulus spatial characteristics on the SIFI remains a pivotal yet unresolved area of research. Two critical questions persist: whether visual stimuli at different locations share a uniform susceptibility to auditory integration, and whether spatial inconsistency diminishes integration levels. While maintaining spatial stability is fundamental to navigating a multisensory environment (Kording et al., 2007), research on auditory spatial cues in SIFI remains sparse due to the auditory channel’s lower spatial resolution compared to vision (Kumpik et al., 2014). Furthermore, evidence regarding the interaction between spatial and temporal characteristics is inconsistent. While most studies suggest an “eccentricity effect”, some studies found no such influence (Gavin et al., 2022; Shulman et al., 1985). Previous paradigms often presented stimuli at randomized locations without spatial cues. Since visual processing efficiency peaks near the central fovea, these studies may have overlooked the influence of unimodal uncertainty and attentional bias on the integration process. Therefore, controlling for spatial attention is essential to isolating the true effect of eccentricity.

This study aims to resolve these discrepancies by systematically investigating the spatial dimensions of audiovisual integration. The first aim of this research examines the effect of visual eccentricity on integration capacity: Experiment 1 extends the parameter range beyond the conventional 10° threshold to map the peripheral visual field more comprehensively, while Experiment 2 utilizes a wider spatial range combined with Bayesian cognitive modeling to characterize the underlying computational mechanisms. The second aim, Experiment 3, explores the role of audiovisual (in)consistency by comparing the effects of ipsilateral, contralateral, and binaural auditory stimuli on illusion perception. By synthesizing behavioral data with computational modeling, this study seeks to delineate how the spatial properties of the visual system and multisensory processing converge into a coherent cognitive mechanism.

## 2. Experiments

### 2.1 Experiment 1: Investigating the Effect of Visual Stimulus Spatial Eccentricity on Audiovisual Integration

This computer-based experiment was designed to replicate the sound-induced flash illusion (SIFI) and, concurrently, explore whether stimuli presented in the central, near-peripheral, and far-peripheral visual fields exhibit differential susceptibility to audiovisual integration. Specifically, we compared the susceptibility of the SIFI across visual stimulus eccentricities of 0°, 7°, and 21°.

For the conditions most likely to induce the SIFI—the 1 Flash 2 Beeps (1F2B) and 2 Flashes 1 Beep (2F1B) conditions—six different levels of Stimulus Onset Asynchrony (SOA) were established: -120 ms, -70 ms, -30 ms, +30 ms, +70 ms, and +120 ms. As illustrated in Figure 1, the sign of the SOA represents the relative temporal order: a negative sign indicates the unimodal stimulus (e.g., flash in 1F2B) preceded the audiovisual pair, while a positive sign indicates it followed. This experiment also served as a preliminary study for parameter refinement during the research process.

**Figure 1.**
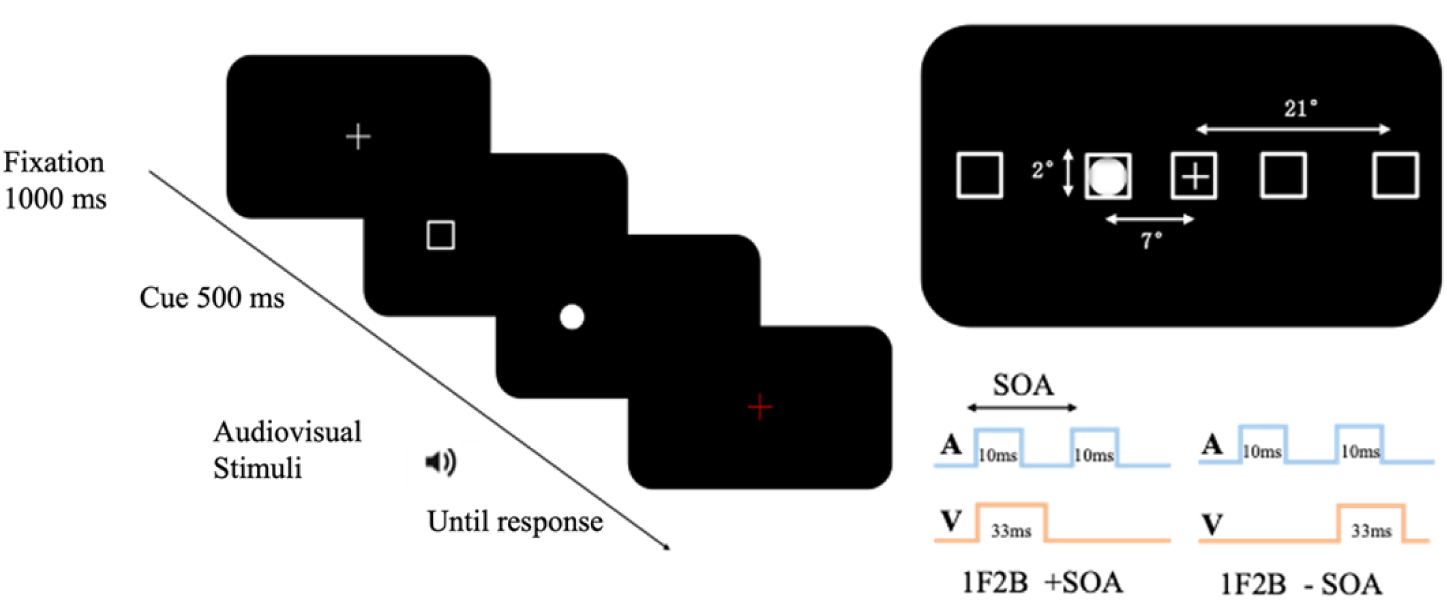
Experimental procedure. Left: task flow of each trial. Visual stimuli are presented at a specific location, while auditory stimuli are delivered binaurally through headphones. Top Right: A schematic showing all possible stimulus locations, example stimuli, and their relative size relationships. In each trial, the cue and the disk (flash) appear at only one specific location. Bottom Right: The temporal sequencesfor a 1F2B (1 Flash, 2 Beeps) trial. This demonstrates the temporal relationship between stimuli under positive and negative SOA. One pair of audiovisual stimuli is always synchronized to begin simultaneously.

Based on previous research on visual stimulus spatial eccentricity and corresponding neuroanatomical evidence (Chen et al., 2017; Falchier et al., 2002; Gavin et al., 2022; Shams et al., 2002; Tremblay et al., 2007), the following hypotheses were formulated: First, the SIFI phenomenon, particularly the fission illusion (where the number of auditory stimuli exceeds the number of flashes), would be reliably replicated across the participant sample, leading to a significant decrease in the correct perception rate. Second, stimuli presented at different spatial locations would exhibit different levels of audiovisual integration; specifically, the 7°and 0°eccentricities (within approximately 10°) would show no significant difference, while the more peripheral 21°eccentricity would be more susceptible to the illusion, consequently yielding a lower reported accuracy rate from participants.

#### Participants

Fifteen university students were recruited. Following preliminary parameter adjustments and accuracy-based screening, nine valid participants (5 female; mean age = 20.44, SD = 1.01) were included in the final analysis. All participants reported normal or corrected-to-normal vision and hearing, were naïve to the purpose of the experiment, and were right-handed. Each participant received a monetary compensation of ¥80 upon completion. All participants provided written informed consent prior to the experiment and received compensation upon completion. The study protocol (with approved No.# 2021-10-18) was approved by the Academic Affairs Committee of the School of Psychological and Cognitive Sciences at Peking University. The above protocol was also applied to the following Experiments 2 and 3.

#### Apparatus and Stimuli

Experiments were conducted in a dimly lit, sound-attenuated laboratory. Visual stimuli were presented on a 27-inch monitor (1920 × 1080 resolution, 144 Hz refresh rate) positioned 60 cm from the participant. The visual target was a white disk (1° radius), and the spatial cue was a white square frame (2° side length). Auditory stimuli (10 ms pure tone, 3000 Hz) were presented via headphones using a sound card with a 96 kHz sampling rate. The experiment was implemented via PsychToolbox-3 (Brainard, 1997; Kleiner M et al., 2007; Pelli, 1997).

#### Experimental Design

The experiment employed a within-subjects design based on the classic SIFI paradigm. The independent variables were stimulus onset asynchrony (SOA, six levels: ±30, ±70, ±120 ms) and visual eccentricity (five levels: -21°, -7°, 0°, 7°, 21°). The dependent variables were response accuracy (proportion of correct flash reports) and reaction time (RT).

#### Experimental Procedure

A strict training protocol ensured task comprehension. Participants completed practice trials with feedback and proceeded to the main experiment only after achieving >90% accuracy. To prevent fatigue, breaks were mandated every 40 trials. As shown in Figure 1, each trial began with a fixation cross (1000 ms), followed by a spatial cue (white frame, 500 ms) appearing at one of the five locations to distinctively guide spatial attention. After a 500 ms gap, the target flash (33 ms) appeared at the cued location. In audiovisual trials, 10 ms beeps (3500 Hz) accompanied the flashes. A red fixation would appear to prompt participants to report the perceived number of flashes via keypress (“Z” or “M”, counterbalanced) within 3 seconds.

Across all trials, excluding the attention check trials (which involved the cue frame but no flash), participants viewed 1 or 2 flashes, accompanied by 0, 1, or 2 auditory stimuli, leading to the 9 combination conditions detailed in Table 1. The core conditions of interest for investigating the SIFI were 1F2B and 2F1B (where F denotes the number of flashes and B denotes the number of beeps), which required the manipulation of the Stimulus Onset Asynchrony (SOA). Specifically, the 1F2B and 2F1B conditions consisted of 12 trials for each combination of the 5 eccentricity levels and 6 SOA levels, totaling 720 trials, which constituted 65% of the total experiment.

**Table 1.** Experiment 1: Trial Distribution.

|  |  | Number of auditory stimuli |  |  |  |
| --- | --- | --- | --- | --- | --- |
|  |  | 0 | 1 | 2 | Sum |
| Number of<br>flashes | 0 | 50 | 15 | 15 | 80 |
|  | 1 | 50 | 100 | 360 | 510 |
|  | 2 | 50 | 360 | 100 | 510 |

The formal experiment comprised a total of 1100 trials, requiring approximately 80 minutes to complete. Participants were instructed to take a minimum one-minute rest after every 50 trials before pressing a key to continue.

#### Data Analysis

The collected response data from all participants were aggregated, and the correct response rates for various conditions were calculated. A repeated-measures Analysis of Variance (ANOVA) was employed for comparisons across levels. It is important to note that the experimental design intentionally oversampled the 1F2B and 2F1B conditions by including more SOA levels, leading to an inherently unbalanced trial distribution.

To ensure a more precise analysis of this data, fully gather evidence supporting all observable effects, and mitigate potential biases arising from the asymmetry between the null and alternative hypotheses (Dienes, 2014), the Bayes Factor analysis method was additionally utilized via JASP software(JASP, 2023). For all Bayesian ANOVAs, the default JASP settings were applied, with the prior r scales for fixed effects, random effects, and covariates set to 0.5, 1 and 0.354, respectively. The interpretation of Bayes Factor values followed the guidelines of Dienes (2014): values greater than 3 represent strong evidence for the alternative hypothesis (*H*_1_); values between 1 and 3 indicate anecdotal support for H1; values between 0.3 and 1 suggest anecdotal support for the null hypothesis (*H*_0_); and values less than 0.3 denote strong evidence for *H*_0_. This approach quantified the relative likelihood of the data under both *H*_0_and *H*_1_, effectively addressing issues related to the unbalanced design and testing biases (Dienes, 2014).

The accuracy rates under different conditions are shown in Figure 2. A two-way repeated-measures ANOVA was conducted on the accuracy rates, with Bonferroni correction applied for post-hoc tests. Both the auditory stimulus and the audiovisual interaction passed Mauchly’s test of sphericity (auditory stimulus: χ² = 2.269, *p* = 0.322; interaction: χ² = 3.361, *p* = 0.186).

**Figure 2.**
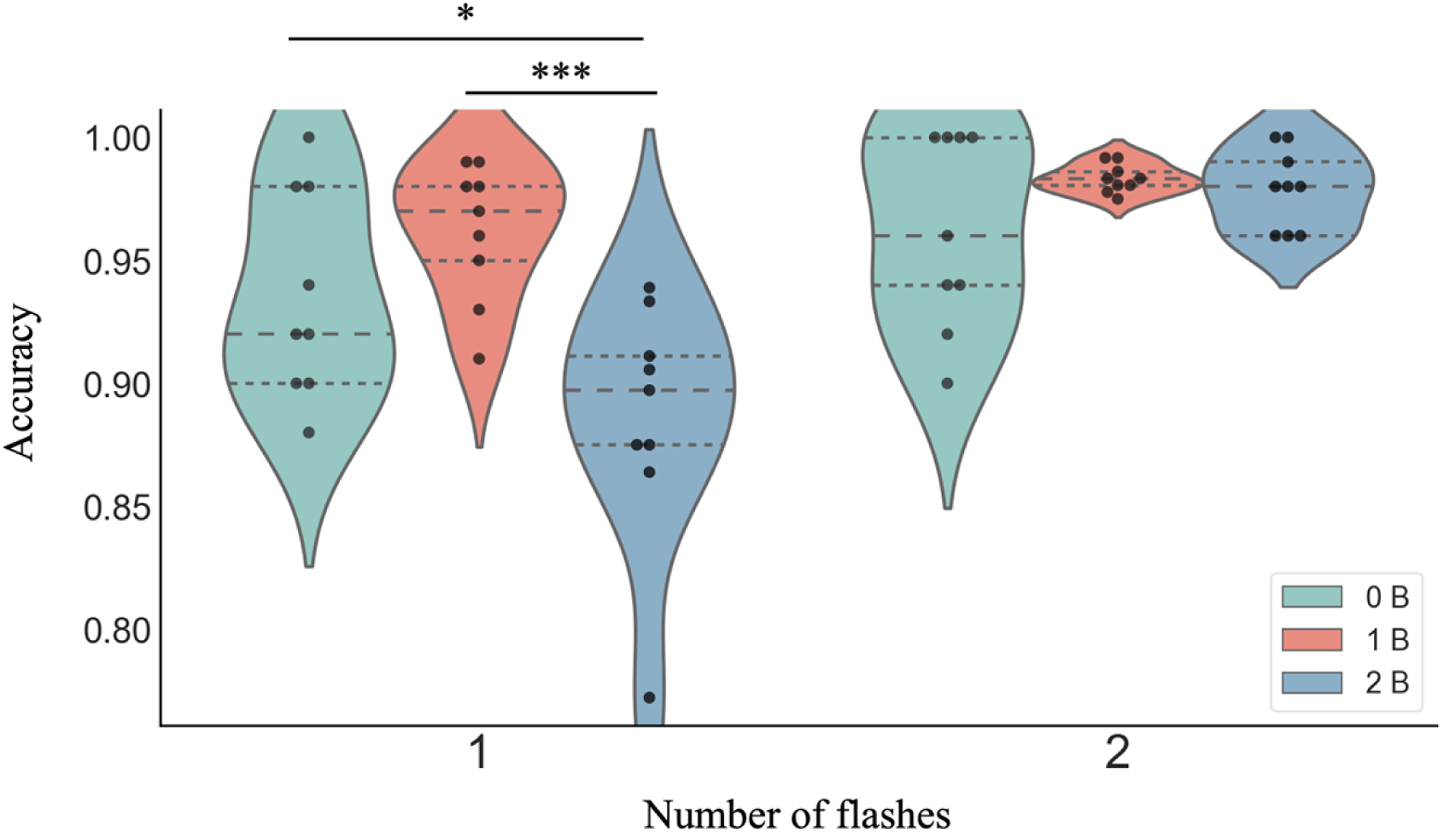
Violin Plots of Participant Report Accuracy Across Conditions in Experiment 1. The width of the violin plot represents the probability density distribution of the data. Each individual dot represents the data point of a single participant under the corresponding condition. The horizontal lines within the violin plots indicate the upper quartile, median, and lower quartile of the data, respectively. * denotes *p*<.05, ** denotes *p*<.01, *** denotes *p*<.001.

The ANOVA results revealed a significant main effect of the number of flashes on participants’ response accuracy, F_(1,8)_ = 14.067, *p* = 0.006, 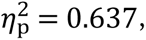 *BF*_10_ = 8.012. Participants’ accuracy in perceiving two flashes was significantly higher, *|MD|* = 0.047, *p* = 0.006, BF₁₀ = 55.529. The number of auditory stimuli also had a significant effect on response accuracy, F_(2,16)_ = 11.421, *p* < 0.001, 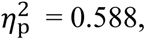 BF₁₀ = 3.704. Under the condition of one auditory stimulus, participants’ response accuracy was significantly higher than with two stimuli (*|MD|* = 0.040, *p* < 0.001, BF₁₀ = 21.489) and with no auditory interference (*|MD|* = 0.024, *p* = 0.038, BF₁₀ = 8.012).

The interaction between the two factors was also significant, F_(2,16)_ = 6.083, *p* = 0.011, 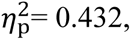 BF₁₀ = 93.434. Simple main effects analysis was conducted, focusing on whether the number of auditory stimuli had different effects under each flash condition. When there was one flash, the simple main effect of sound stimuli was significant, F_(2,16)_ = 20.792, *p* < 0.001, BF₁₀ = 83.776. The difference between 0B and 1B was not significant, *|MD|* = 0.027, *p* = 1.000, BF₁₀ = 0.900. However, accuracy under 0B was significantly higher than under 2B, *|MD|* = 0.050, *p* = 0.024, BF₁₀ = 1.985. Accuracy under 1B was also significantly higher than under 2B, *|MD|* = 0.076, *p* < 0.001, BF₁₀ = 111.291. This indicates a clear flash fission illusion: when the number of sound stimuli exceeded the actual number of flashes, participants’ subjective reports of the number of flashes also increased.

When there were two flashes, the number of auditory stimuli had no significant effect on accuracy, F_(2,16)_ = 1.712, *p* = 0.212, BF₁₀ = 0.928. Under two flashes, participants’ reported accuracy was high across different numbers of auditory stimuli, and no significant flash fusion illusion was observed.

Focusing further on participants’ accuracy when a single flash was presented at different eccentricities, we conducted a two-way repeated-measures ANOVA with factors of eccentricity and number of auditory stimuli. The interaction between eccentricity and auditory number violated sphericity (χ²₃₅ = 100.573, *p* < 0.001), so degrees of freedom were adjusted with the Greenhouse–Geisser correction.

As shown in Figure 3, the main effect of spatial eccentricity was significant, F_(4,32)_= 9.342, *p* < 0.001, 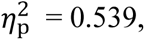 BF₁₀ = 85.524. Accuracy at 21° in both hemifields was lower than in central vision (left 21°: *|MD|* = 0.055, *p* = 0.004, BF₁₀ = 170.112; right 21°: *|MD|* = 0.070, *p* < 0.001, BF₁₀ = 489.405). In addition, accuracy differed between 7° and 21° in both hemifields (left: |*MD|* = 0.045, *p* = 0.028, BF₁₀ = 9.183; right: *|MD|* = 0.049, *p* = 0.013, BF₁₀ = 10.061), whereas performance at 7° did not differ from central vision. The main effect of auditory-stimulus number was also significant, F_(2,16)_ = 10.622, *p* = 0.001, 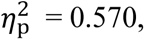 BF₁₀ = 29.283. The eccentricity × auditory-number interaction was not significant after correction, F_(3.504,28.035)_ = 1.583, *p* = 0.211, 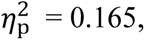 BF₁₀ = 0.882.

**Figure 3.**
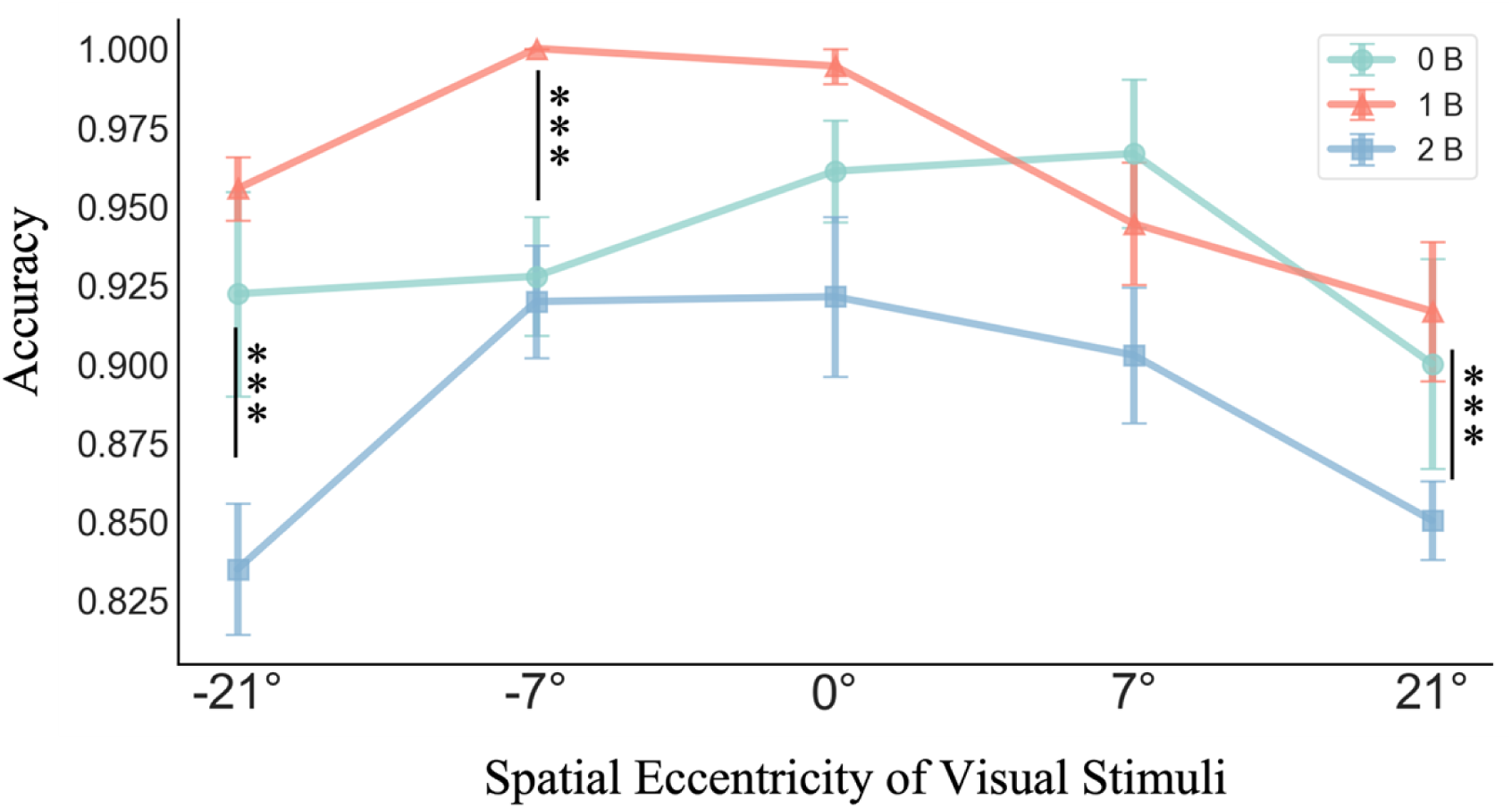
Response accuracy across conditions for trials with a single flash in Experiment 1. Different colored lines represent different numbers of auditory stimuli. Error bars indicate one standard error (SE). * denotes *p*<.05, ** denotes *p* <.01, *** denotes *p* <.001.

Simple-main-effect analyses examined whether the eccentricity profile was equivalent across auditory conditions. Without auditory distractors, eccentricity had no significant impact on accuracy, F_(4,32)_ = 1.332, *p* = 0.280, BF₁₀ = 0.330. In contrast, when one or two auditory stimuli were presented, eccentricity strongly modulated accuracy (1B: F_(4,32)_= 8.630, *p* < 0.001, BF₁₀ = 623.599; 2B: F_(4,32)_= 10.217, *p* < 0.001, BF₁₀ = 1 093.603). Thus, no reliable eccentricity effect emerged in the unimodal visual task, whereas introducing auditory stimuli in a cross-modal setting revealed marked performance differences between peripheral and peri-foveal locations.

Prior analyses confirmed the presence of sound-induced flash illusions and showed that the spatial position of visual stimuli modulates audiovisual integration. Temporal alignment is also critical, as the inter-stimulus interval systematically shapes susceptibility to the SIFI. We therefore investigated whether the impact of spatial eccentricity varies across temporal contexts—specifically, whether a space–time interaction exists.

Figure 4 illustrates performance in fission-illusion trials (F1B2) as a function of SOA and eccentricity.

**Figure 4.**
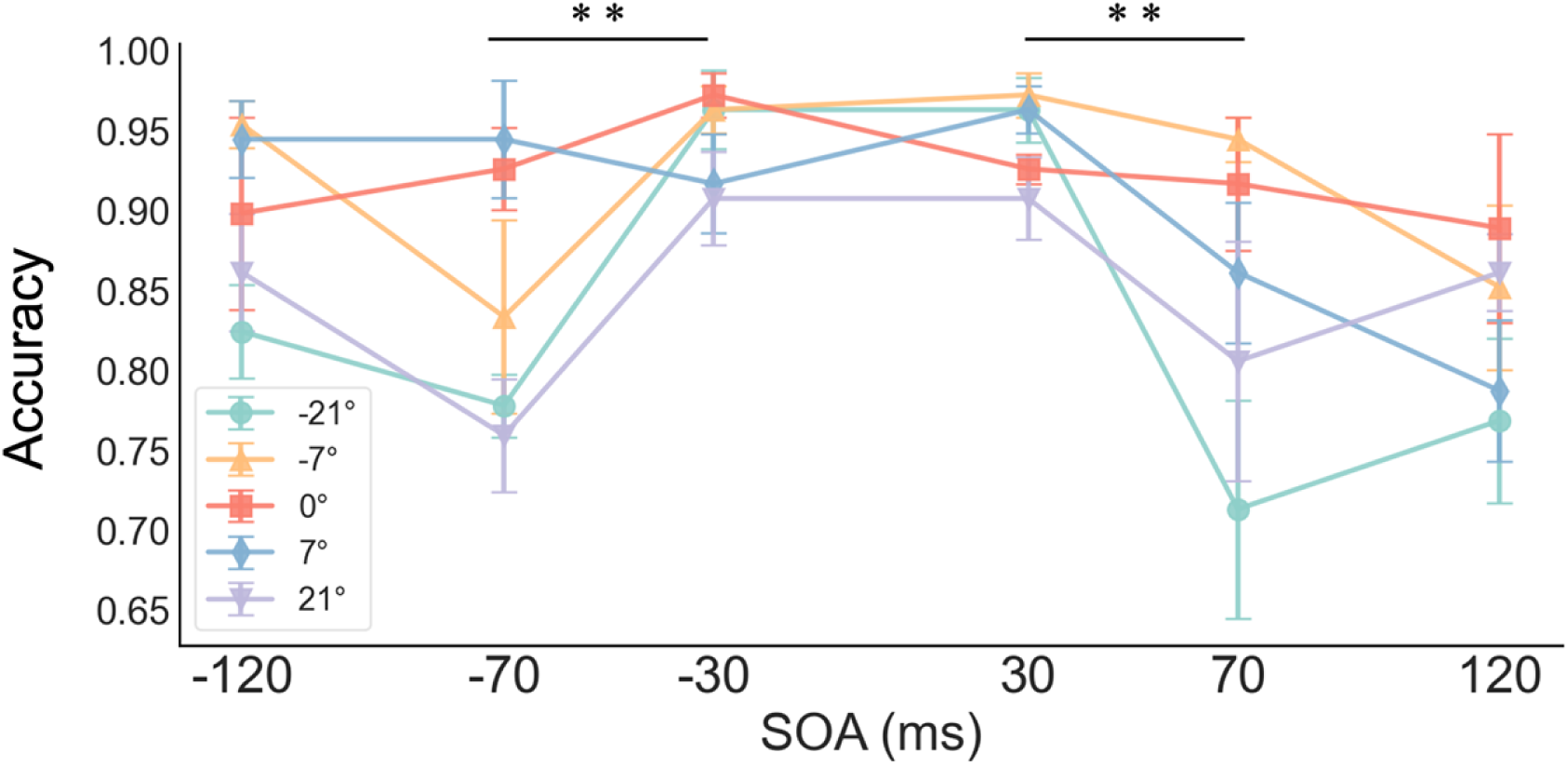
Response accuracy in the fission illusion condition (1F2B) across different SOAs. Different colored lines represent the various spatial locations of the visual stimuli. Error bars indicate one standard error (SE). * denotes *p* <.05, ** denotes *p* <.01, *** denotes *p* <.001

A two-way repeated-measures ANOVA (eccentricity × SOA) revealed significant main effects of both eccentricity, F_(4,32)_ = 10.217, *p*< 0.001, 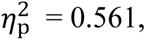 BF₁₀ = 17.954, replicating earlier findings, and SOA, F_(5,40)_ = 8.078, *p*< 0.001, ηp² = 0.502, BF₁₀ = 182.648. Contrary to the expectation that shorter SOAs should promote stronger integration, accuracy at |SOA| = 30 ms was significantly higher than at |SOA| = 70 ms (*p*< 0.01), with no other pairwise SOA comparisons reaching significance. Additionally, a significant eccentricity × SOA interaction emerged, F_(20,160)_ = 2.233, *p* = 0.003, 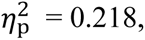 BF₁₀ = 67.643.

Simple-main-effect analyses revealed that eccentricity reliably influenced performance exclusively within the SOA = –70 ms and +70 ms windows (–70 ms: F_(4,32)_ = 5.372, *p* = 0.002, BF₁₀ = 42.601; +70 ms: F_(4,32)_ = 4.368, *p* = 0.006, BF₁₀ = 10.316). These are precisely the SOAs that maximized illusion susceptibility. Thus, when auditory and visual signals are temporally discrepant yet still integrated, the spatial location of the visual event determines the strength of that integration. At SOAs that are too brief or too prolonged—conditions in which observers appear largely immune to auditory influence—the modulatory effect of spatial position disappears.

#### Discussion

We successfully replicated the fission illusion (Shams et al., 2002) and extended the findings to the spatial domain. Our results demonstrate that while SIFI susceptibility is stable within the central 10°, it significantly increases in the far periphery (21°). Importantly, by using pre-cues to equate spatial attention, we ruled out the possibility that this effect stems from reduced peripheral attention or visual acuity.

These findings partially align with earlier work showing stronger SIFI at peripheral locations (Chen et al., 2017; Shams et al., 2002; Tremblay et al., 2007). First, previous studies sampled only ≤ 10° eccentricity and reported marginal or null differences; we likewise found no change between 0° and 7°, consistent with Gavin et al. (2022). Second, by extending the spatial span to 21°, we reveal a steep increase in illusion susceptibility, while unimodal visual sensitivity remains unchanged. Audiovisual integration therefore exhibits a distinctive spatial signature, suggesting that visual input from different spatial locations deploy different integration strategies or weighting schemes.

Our results deviate from the assumption that decreasing SOAs monotonically increase illusion strength. At ±30 ms, accuracy was significantly high—exceeding baseline levels—suggesting facilitation over fission. This suggests that when auditory stimuli are too close in time, they may be perceptually fused or fall within a single cycle of neural oscillation, failing to trigger the “two-beep” induced fission (Fiebelkorn & Kastner, 2019).

Taken together, the results broaden the known spatial landscape of the SIFI. But we still cannot adjudicate between two mechanistic accounts: (a) greater visual uncertainty in the periphery biases the brain’s optimal estimate toward the auditory count (Shams & Beierholm, 2010), and (b) stronger direct connectivity between auditory cortex and the peripheral representation of early visual cortex (Eckert et al., 2008) gives auditory input heavier weight. To further disentangle whether this spatial effect arises from sensory uncertainty or integration weights, Experiment 2 will expand the eccentricity range and employ Bayesian modeling.

### 2.2 Experiment 2: Extended Spatial-Eccentricity for SIFI with Hierarchical Bayesian Modelling

Building on the previous findings, we exploited a 360° acoustic arena to sample observer performance in the SIFI paradigm at five eccentricities (0°, 15°, 30°, 45°, 60°; 15° steps). Using hierarchical Bayesian modelling anchored in the classical causal-inference framework (Shams et al., 2006; Shams & Beierholm, 2010), we compared two families of models: (1) a “visual-uncertainty” family that assumes fixed AVI weights but allows visual likelihood variance to increase with eccentricity, and (2) an “AV-weight” family that keeps likelihood variance constant while letting the prior weight assigned to the common-cause hypothesis vary with retinal location.

At the behavioral level, we predict that across the 0–60° eccentricity range, visual accuracy will decline and reports will become increasingly biased by the number of auditory beeps, while performance on unimodal (flash-only) and congruent audiovisual trials remains invariant. Computationally, if the uncertainty model family provides a superior fit, it would favor the classic view that audiovisual integration (AVI) computations are spatially identical, with performance deficits driven solely by increased peripheral visual noise; however, a superior fit for the AV-weight family would align with neuroanatomical evidence (Eckert et al., 2008; Falchier et al., 2002; Rockland & Ojima, 2003), suggesting that different retinotopic loci possess intrinsic susceptibilities to auditory influence, modeled as location-specific prior weights within a Bayesian causal-inference framework.

#### Participants

Thirteen undergraduate students took part in the experiment. After applying an accuracy criterion, data from eleven participants (six female) were retained.

Participants’ ages were ranged from 19 to 22 years (M = 20.27, SD = 1.01). All participants had normal hearing and normal or corrected-to-normal vision, were right-handed, and had no prior experience with similar experiments. Each participant received 80 RMB in cash after the session.

#### Apparatus and stimuli

The experiment was conducted in a single, well-ventilated laboratory under dim ambient lighting. As shown in Figure 5 (left), participants performed the computer-based task on a 1.4-m-radius curved “audio-screen” display (resolution 1920 × 1080, refresh rate 60 Hz). Sounds were delivered via a sound card sampled at 44.1 kHz and presented through closed-back monitor headphones worn throughout the experiment.

**Figure 5.**
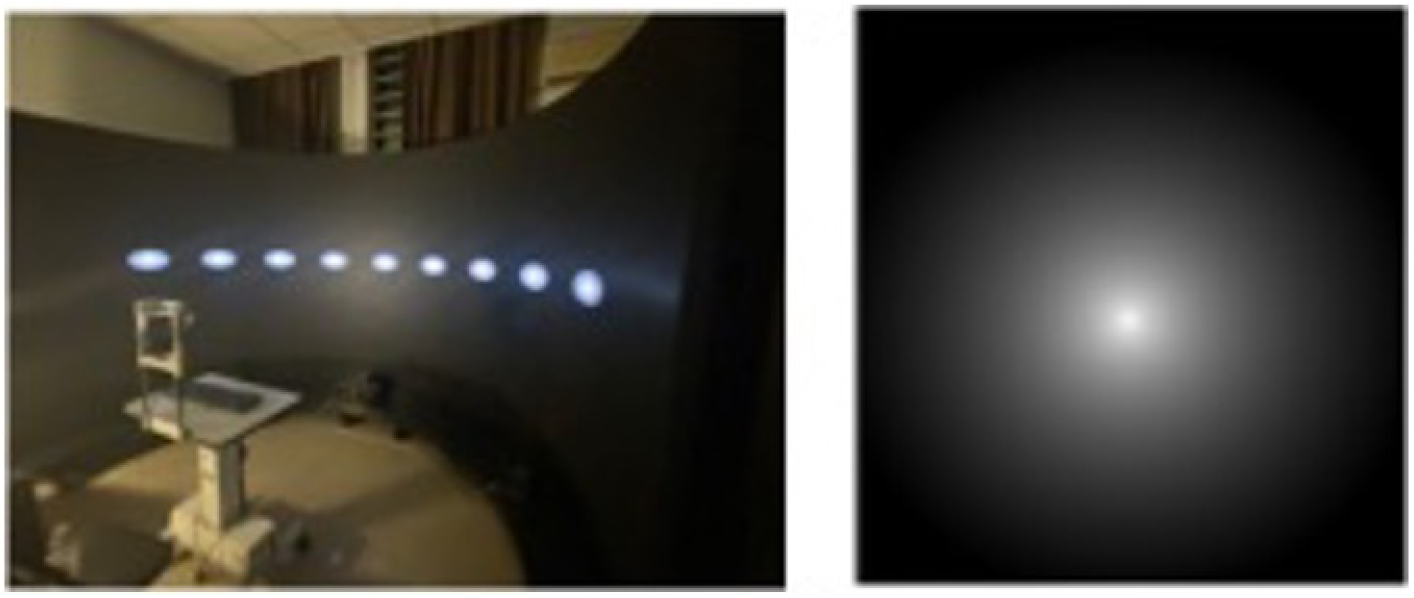
Experimental Scene and Stimulus Examples for Experiment 2. Left: An illustration of the laboratory setting. The screen displays all possible locations for stimulus presentation; however, during an actual trial, the flash appears at only one specific location. Right: The gradient disk stimuli used in Experiments 2 and 3, which feature logarithmic decay along the axial direction.

Visual stimuli were white, Gaussian-ramped disks (Figure 5, right) chosen to minimise sharp-edge after-effects (Stiles et al., 2020). A white square frame (4° × 4°) served as the location cue. With the viewing distance fixed at 1.4 m by a chin-rest, the disk subtended 2° of visual angle. Auditory stimuli were 10-ms pure tones at 2000 Hz. Participants responded using three keyboard keys (“Z”, “?/” and spacebar) while maintaining a stable head position.

#### Design and Procedure

As in Experiment 1, flashes were delivered at different spatial locations while the number of accompanying beeps varied. The critical independent variable was visual eccentricity, with nine levels: −60°, −45°, −30°, −15°, 0°, 15°, 30°, 45°, and 60°. Because we focused on spatial rather than temporal properties of audiovisual integration, SOA was not factorially manipulated; instead, only two SOAs—40 ms (“short”) and 70 ms (“long”)—were used for the single-channel continuation stimuli in both fission (more beeps than flashes) and fusion (more flashes than beeps) blocks. Whenever both modalities were stimulated, the first audiovisual pair was always presented simultaneously; subsequent unimodal stimuli followed at the designated SOA.

The stimulus set was expanded (0-3 flashes; 0-2 beeps) to increase difficulty and discourage response bias. Three-flash trials were used as a quality control measure, with a 50% accuracy threshold for participant exclusion. As shown in Table 2, the primary fission and fusion conditions consisted of 12 trials per SOA and eccentricity level. These were embedded within a total of 1,150 trials. To manage fatigue, participants took mandatory breaks for at least one minute every 80 trials. Total experimental duration was approximately 90 minutes.

**Table 2.** Experiment 2: Trial Distribution.

|  |  | Number of auditory stimuli |  |  |  |
| --- | --- | --- | --- | --- | --- |
|  |  | 0 | 1 | 2 | Sum |
| **Number of**<br>**Flash** | 0 | (No.of auditory stimuli is random) |  |  | 43 |
|  | 1 | 90 | 135 | 216 | 441 |
|  | 2 | 90 | 216 | 135 | 441 |
|  | **3** | **45** | **90** | **90** | **225** |

#### Procedure

The task closely followed Experiment 1. After stimulus offset, an “X” appeared at the bottom of the screen to signal the response window; the deadline was shortened to 2.5 s and the inter-trial interval to 0.75 s to reduce overall duration. Participants pressed “Z” or “?/” to report “1” or “2” flashes (key mapping counter-balanced across subjects); the space-bar was used for trials containing three flashes. All other procedural details were identical to Experiment 1.

#### Data analysis

Responses were pooled and the mean reported number of flashes calculated for each condition. Repeated-measures ANOVAs were used for factorial comparisons. To mitigate the imbalance in trial counts, Bayesian factors were again computed with JASP (JASP Team, 2023).

Figure 6 shows the mean reported flashes.

**Figure 6.**
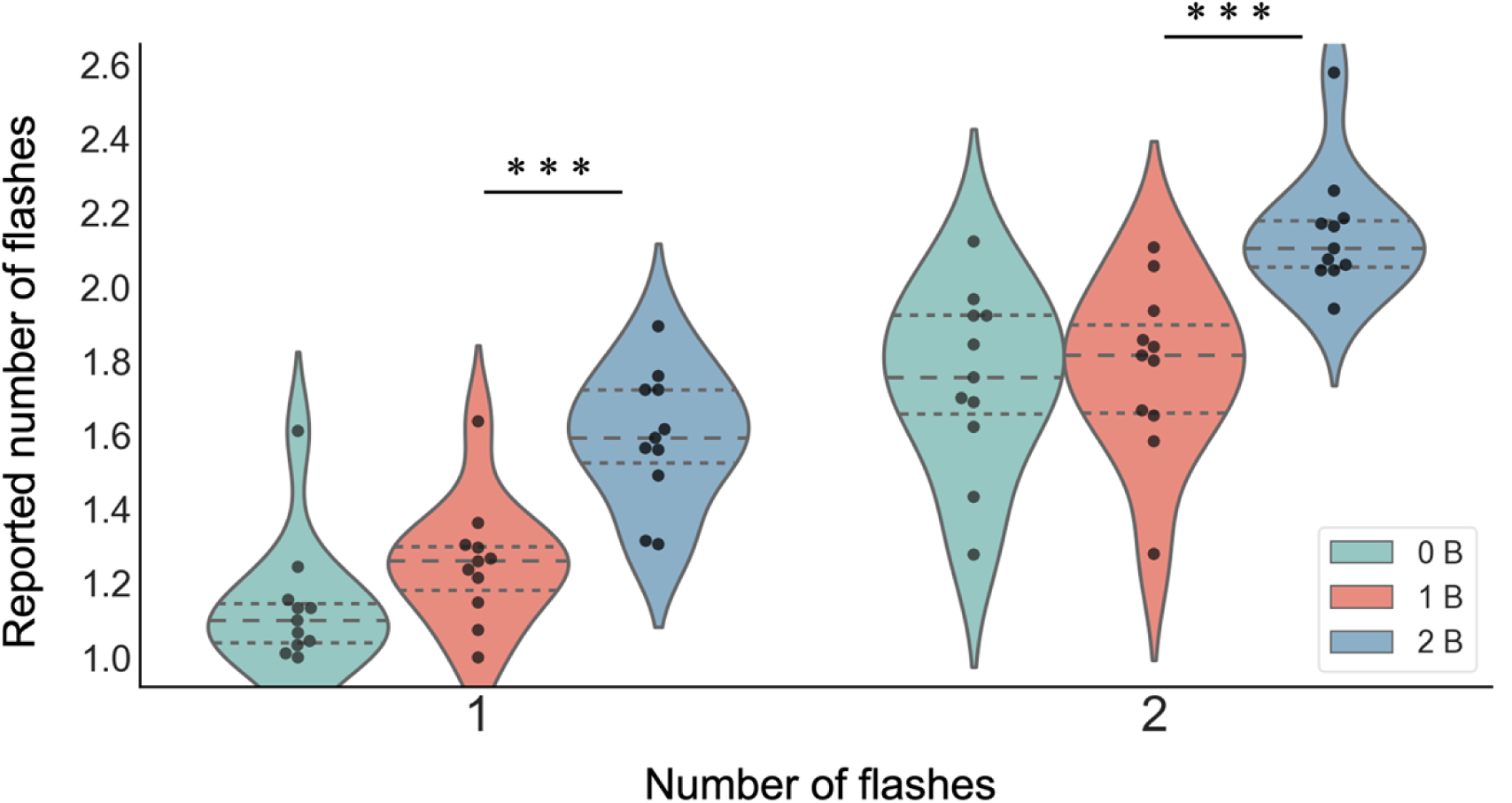
Violin Plots of Participant Response Accuracy Across Conditions in Experiment 2. The width of each violin plot represents the probability density distribution of the data. Individual dots represent data points for each participant within that specific condition. The horizontal lines within the violin plots denote the upper quartile, median, and lower quartile of the data. * p<.05, ** p<.01, *** p<.001

A 2 (target flashes: 1 vs. 2) × 3 (auditory beeps: 0, 1, 2) within-subjects ANOVA analysis revealed significant main effects of both flashes and beeps, with no interaction between them. For flashes, participants reported significantly more flashes when two were presented compared to one (F(1, 10) = 156.11, p < .001, ηp² = .940, BF₁₀ = 1.24 × 10⁵), with a mean difference of 0.563 (p < .001, BF₁₀ = 2.20 × 10¹⁴).

For beeps, there was a significant main effect (F(2, 20) = 42.11, p < .001, ηp² = .808, BF₁₀ = 2.41 × 10⁵), where two beeps increased flash reports relative to both zero beeps (|MD| = 0.426, p < .001, BF₁₀ = 5.29 × 10⁶) and one beep (|MD| = 0.353, p < .001, BF₁₀ = 5.33 × 10⁴). However, the interaction between flashes and beeps was not significant (F(2, 20) = 0.83, p = .450, ηp² = .077, BF₁₀ = 0.321), indicating that the effect of flashes was consistent across beep conditions.

Bonferroni-corrected post-hoc tests (Figure 6) confirmed the fission illusion: for 1-flash trials, 0B ≈ 1B (|MD| = 0.115, p = .946, BF₁₀ = 8.66), but 2B > 1B > 0B (all ps < .001). For 2-flash trials, 2B again produced higher reports than 1B or 0B (|MD|s ≥ 0.367, ps < .001). No reliable fusion illusion was observed; instead, two beeps generally increased flash reports, consistent with cross-modal summation. All subsequent analyses therefore focus on 1-flash trials to characterise the spatial profile of fission.

Focusing again on 1-flash trials, we submitted accuracy to a 2-way repeated-measures ANOVA with factors Auditory Level (0B, 1B, 2B) and Eccentricity (−60° to +60° in 15° steps). Both factors passed Mauchly’s test (auditory: χ²₂ = 5.296, p = .071; eccentricity: χ²₃₅ = 33.423, p = .663).

The results demonstrated significant main effects of auditory beeps and eccentricity, as well as a significant interaction between them. For auditory level, the characteristic fission pattern is dominant (F(2, 20) = 38.42, p < .001, ηp² = .367, BF₁₀ = 7.99 × 10⁴), with two beeps eliciting more flash reports than both one beep and zero beeps (all ps < .001), which did not differ significantly. Eccentricity also significantly affected reports (F(8, 80) = 8.86, p < .001, ηp² = .112, BF₁₀ = 4.10 × 10⁴): central vision (0°) yielded lower (more accurate) reports than every peripheral location (ps < .05), and right 15° produced lower reports than right 45° (|MD| = 0.214, p = .006, BF₁₀ = 71.16 (Figure 7). Critically, the auditory × eccentricity interaction was significant (F(16, 160) = 3.17, p < .001, ηp² = .072, BF₁₀ = 1.43 × 10³). Simple-main-effect analyses revealed that eccentricity had no effect in the unimodal visual condition (0B: F(8, 80) = 1.52, p = .163, BF₁₀ = 0.37) and only a marginal effect in the congruent audiovisual condition (1B: F(8, 80) = 1.96, p = .062, BF₁₀ = 0.999). However, in the conflict condition (2B), a strong eccentricity effect was observed (F(8, 80) = 18.18, p < .001, BF₁₀ = 5.39 × 10⁷), where reports approached the veridical count of one flash only in central vision, while all peripheral locations showed significantly more reports of flashes, indicating stronger fission illusions in the visual periphery.

**Figure 7.**
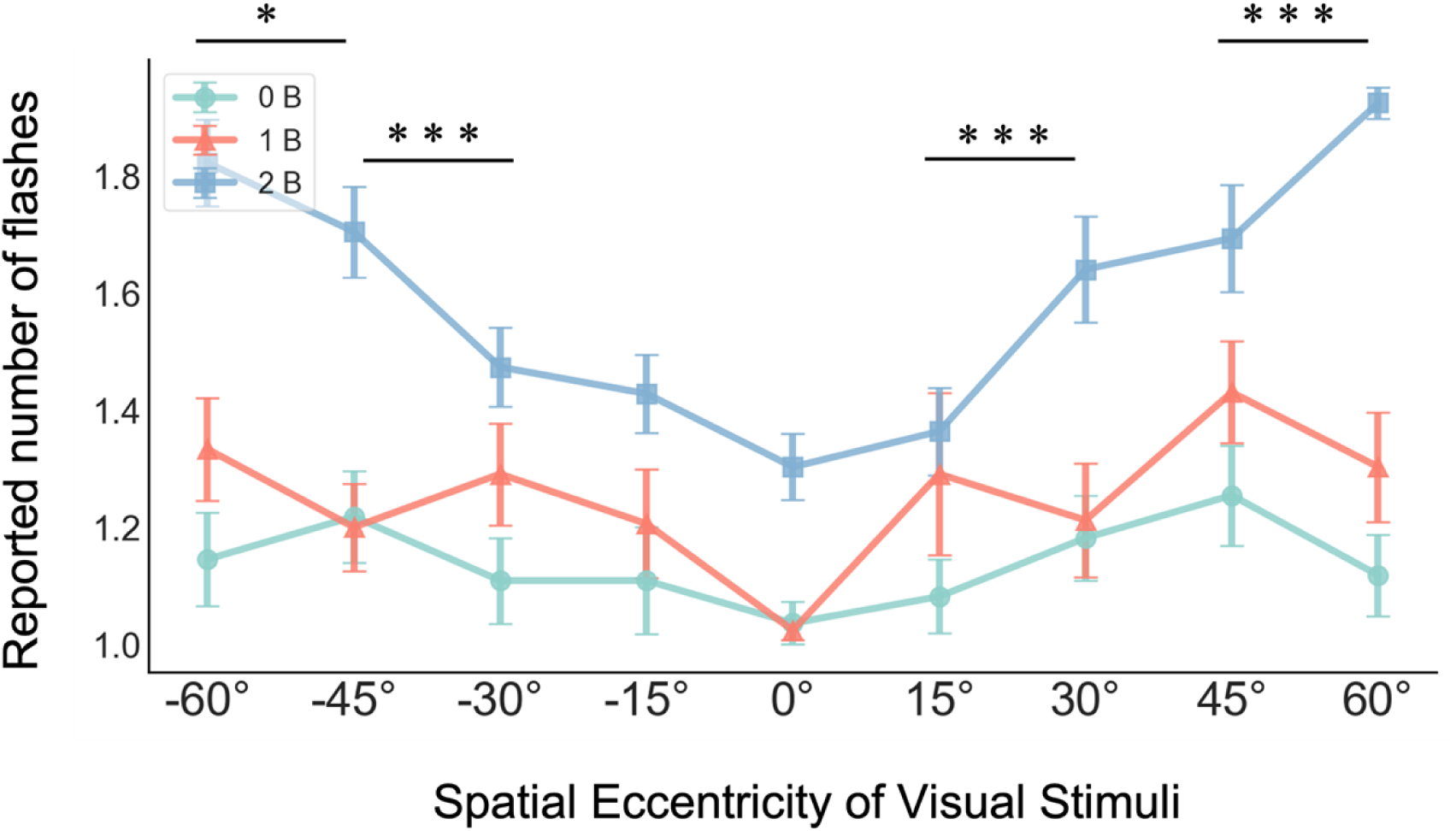
Mean reported number of stimuli for single target-flash trials across auditory conditions in Experiment 2. Different colored lines represent different numbers of auditory stimuli. Error bars indicate one standard error of the mean (SEM). * denotes p<.05, ** denotes p<.01, *** denotes p<.001.

Together with Experiment 1, these findings confirm that when attention is largely equated across the visual field, unisensory visual perception is spatially flat, whereas multisensory processing—especially under audiovisual conflict—is sharply modulated by retinal eccentricity.

In the 1F2B condition, we used two SOAs: 40 ms and 70 ms. To test whether spatial eccentricity interacts with temporal context, we compared performance across eccentricity and SOA. Because the target was always one flash and the distractors always two beeps, accuracy and reported count are perfectly inversely related, and Figure 8 shows that accuracy follows an inverted Gaussian profile across space. We therefore used hit-rate as the metric and assumed a Gaussian relationship between eccentricity x and P(hit):

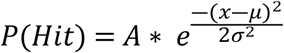

**Figure 8.**
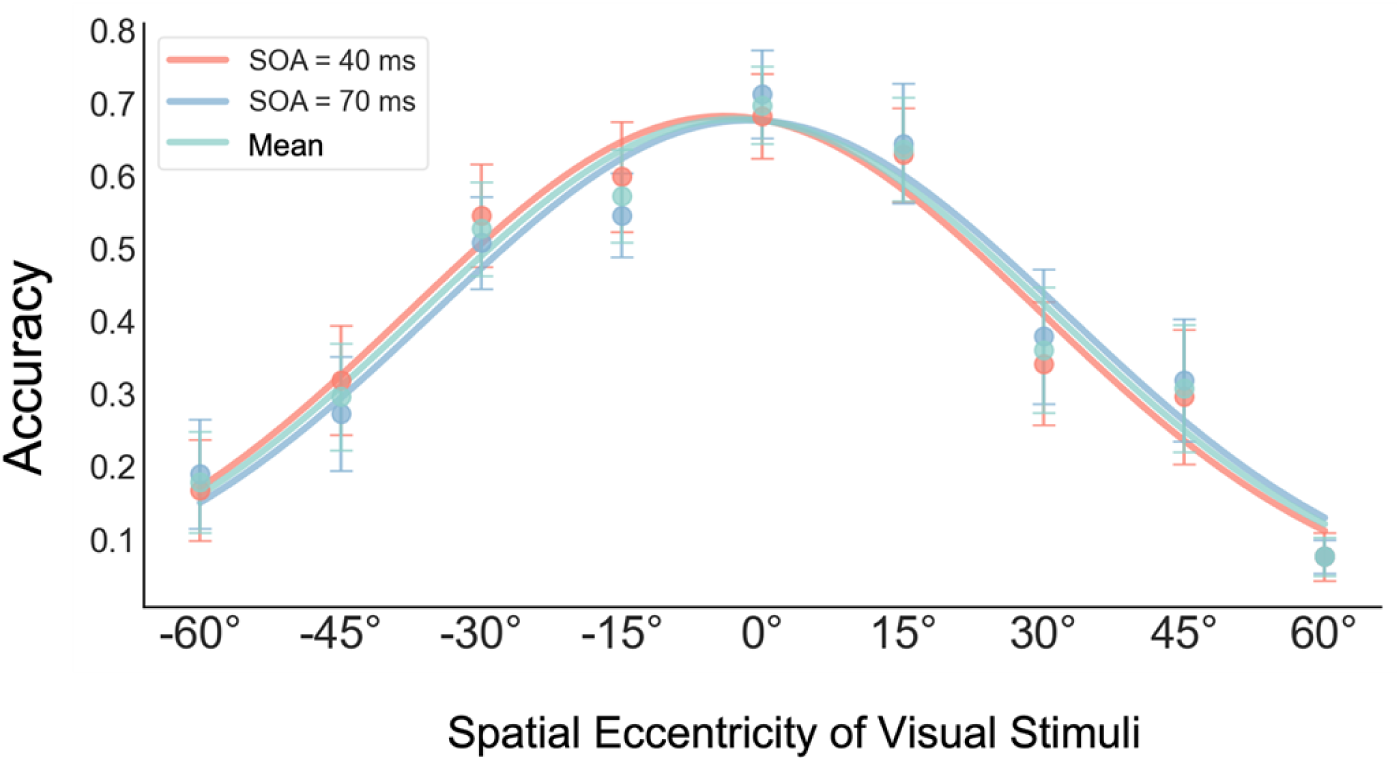
Participant response accuracy across eccentricities in the 1F2B condition of Experiment 2. Different colored lines represent various SOA conditions and the grand average of the data; error bars indicate one standard error (SE).

Data for each SOA were fitted separately; the two SOAs were also collapsed to obtain an average curve. Adjusted R² was computed for each model. Table 3 summarises the parameters. In all cases, the Gaussian described the data well (adjusted R² > .94) and the curves for 40 ms and 70 ms were almost superimposable. Thus, univariate accuracy can be characterised by a central-peaked Gaussian that declines toward the periphery. What remains unknown is whether this spatial gradient reflects auditory interference or merely weaker unisensory vision in the periphery; the modelling analyses that follow will address this question.

**Table 3.**
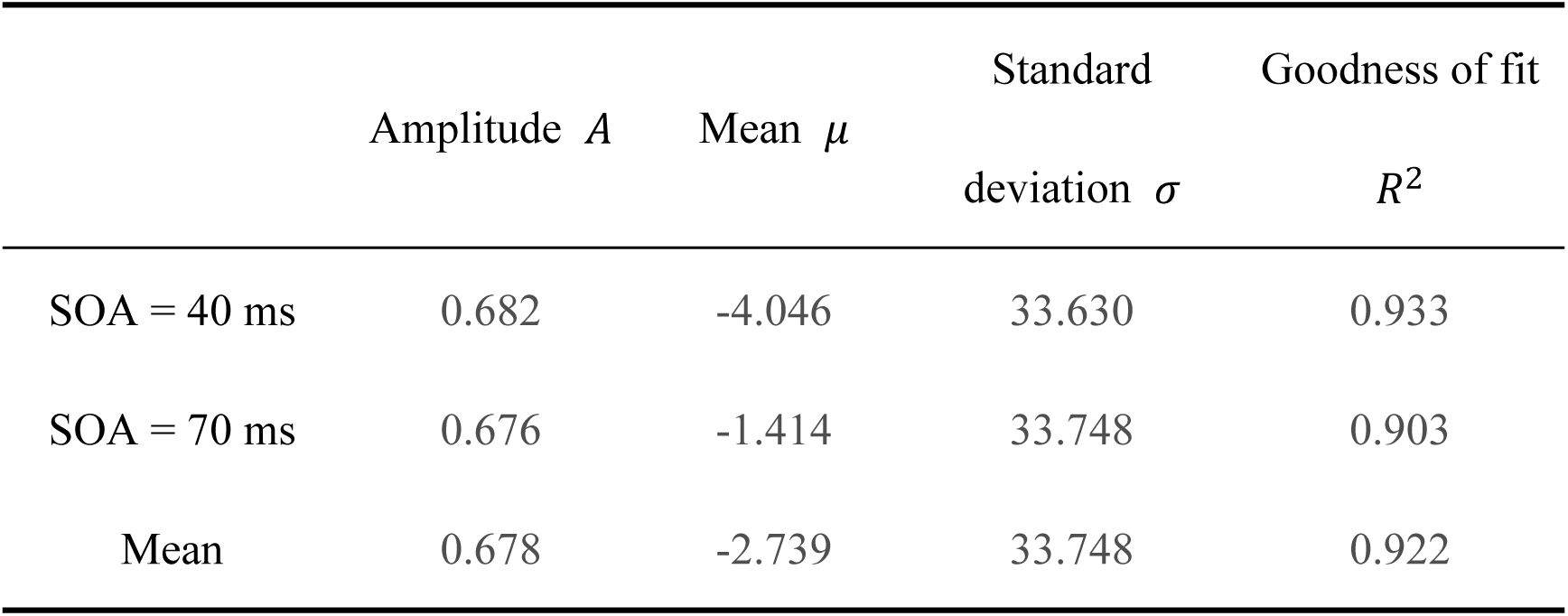
Table of Gaussian Curve Fitting Parameters for Response Accuracy in 1F2B Trials.

| | Amplitude $A$ | Mean $\mu$ | Standard<br>deviation $\sigma$ | Goodness of fit<br>$R^2$ |
| --- | --- | --- | --- | --- |
| SOA = 40 ms | 0.682 | -4.046 | 33.630 | 0.933 |
| SOA = 70 ms | 0.676 | -1.414 | 33.748 | 0.903 |
| Mean | 0.678 | -2.739 | 33.748 | 0.922 |

The hit-rates for the two SOA conditions, together with the collapsed data and their fitted Gaussian curves, are plotted in Figure 8. Accuracy was virtually identical at 40 ms and 70 ms, with no discernible separation. A 9 (eccentricity) × 2 (SOA) repeated-measures ANOVA confirmed a significant main effect of eccentricity, F(8, 80) = 18.18, p < .001, ηp² = .645, BF₁₀ = 1.73 × 10¹², replicating the previous analysis, but neither a significant main effect of SOA, F(1, 10) = 0.14, p = .895, ηp² < .001, BF₁₀ = 0.24, nor a significant eccentricity × SOA interaction, F(8, 80) = 0.72, p= .671, ηp² = .007, BF₁₀ = 0.09.

Thus, within the two SOAs tested, audiovisual integration was unaffected by temporal separation. Given we focuses on spatial rather than temporal characteristics, the 40-ms and 70-ms data were pooled for all subsequent modelling analyses.

#### Bayesian modelling

The findings from Experiments 1 and 2 demonstrate that the spatial location of visual stimuli significantly modulates SIFI perception, suggesting that different regions of the visual field may utilize distinct audiovisual integration mechanisms. Integrating modality reliability theory with the modeling framework established by Hirst et al. (2020)(Hirst et al., 2020), we propose two hypotheses to account for this spatial variation in multisensory capacity.

First, *The Spatial Weighting Hypothesis.* The direct cross-modal influence of auditory stimulation may have a greater impact in the peripheral visual field, resulting in greater informational weight being assigned to the auditory modality during the integration process. Combined with responses that integrate auditory information, this makes illusory perception more likely to occur.

Second, *the Visual Uncertainty Hypothesis*: This account posits that the increase in illusory percepts in the periphery is a direct consequence of reduced visual reliability. As visual acuity declines with eccentricity, the uncertainty surrounding visual numerosity perception increases. Within a Bayesian framework, the brain compensates for this unreliable visual signal by relying more heavily on the relatively more precise auditory information, leading to the perception of the sound-induced flash illusion.

To investigate which mechanism better supports the current findings, we implemented Bayesian modeling to perform hierarchical inference about internal cognitive processes (Van De Schoot et al., 2021). We used PyMC5, a Python library for probabilistic programming that offers extensive choices of prior and posterior distributions, for model construction and comparison, and implemented algorithms such as Markov Chain Monte Carlo (MCMC) for posterior sampling (Abril-Pla et al., 2023).

Following the Bayesian ideal observer model (Shams & Beierholm, 2010), we model perceived numerosity as a weighted combination of auditory and visual information. Sensory inputs are treated as probability distributions; when signals originate from the same source, their likelihoods are multiplied to create a joint audiovisual representation. The observer then forms a final perceptual estimate by calculating the precision-weighted average of the individual sensory channels and the integrated likelihood. This approach ensures that the resulting percept is a reliable inference based on the relative uncertainty of each modality.

Mathematically, this process can be described as follows: when an observer forms a perceptual representation *S̄*_*v*_ of visual information based on received audiovisual sensory information x_*v*_ and x_a_, the following occurs:

First, if the observer believes that visual and auditory stimuli originate from different sources (C = 2, where C represents causal structure, i.e., the number of sources), and can independently receive information from both modalities, the sensory information received through each modality is, due to the presence of noise, essentially represented as Gaussian distributions. The means *μ*_*v*_ and *μ*_*a*_represent individual subjective estimates, while standard deviations *σ*_v_ and *σ*_a_ represent the uncertainty of unisensory information:

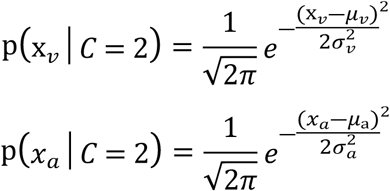

When an individual cannot completely separate auditory and visual stimuli in perception, integration occurs. In this case, the observer believes that the audiovisual stimuli originate from the same source (C = 1), and subsequently forms a unified representational estimate of the audiovisual stimuli. Computationally, this audiovisual representation x_av_ is obtained by multiplying the likelihood functions of the unisensory auditory and visual channels, and the result is also a Gaussian distribution:

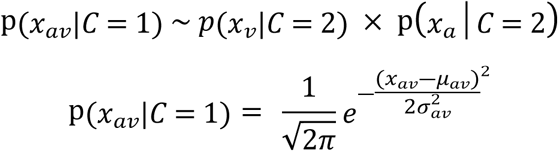

Here, both the mean *μ*_*av*_ and standard deviation *σ*_av_ of the audiovisual stimulus estimate can be expressed in terms of the distribution parameters of the unisensory stimuli:

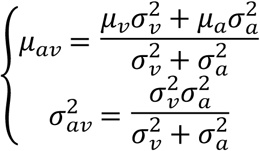

When making perceptual decisions, according to Bayes’ formula, individuals will form weighted averages of the unisensory inference *S̄* _*v*,*C*=2_ and the integrated inference *S̄* _a*v*_ using the probabilities of same-source versus different-source scenarios as weights, thereby generating the optimal inference for unisensory information.

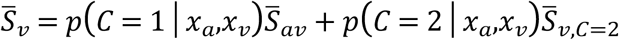

Using weight (w) to represent the probability of audiovisual integration occurring in subjects, we obtain:

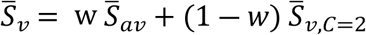

This hierarchical inference structure can be well characterized by Bayesian models, and through manipulation of different parameters, we can investigate the source of differences in subjects’ audiovisual integration levels across various spatial eccentricities. To explore the specific mechanisms underlying these spatial distribution characteristics, we constructed five models as shown in Figure 9. The models share these common variables: *v* (visual information likelihood, representing the observer’s sensory estimate of visual information) and *a* (auditory information likelihood, representing the sensory estimate of auditory information), which are sampled from two Gaussian distributions. Since this involves fitting the 1F2B condition, sampling is performed from distributions with a mean of 1 and standard deviation of *σ*_*v*_ and a mean of 2 and standard deviation of *σ*_*a*_, respectively. The standard deviation parameters *σ*_*v*_ and *σ*_*a*_, which characterize the uncertainty of unisensory information, are free parameters in the model that need to be fitted through data sampling. Observers combine noisy audiovisual signals and their respective uncertainties to form a unified stimulus representation, assuming the signals originate from the same source (*av*). Since the final weighted average is essentially a weighted average of the means, only *μ*_*av*_ needs to be calculated during the sampling process. Finally, the observer’s estimate of visual perception is obtained through weighted averaging of *μ*_*av*_ and their own sensory sample *v* to produce the optimal estimate (opt), where the weight *w* is also a free parameter of the model.

**Figure 9.**
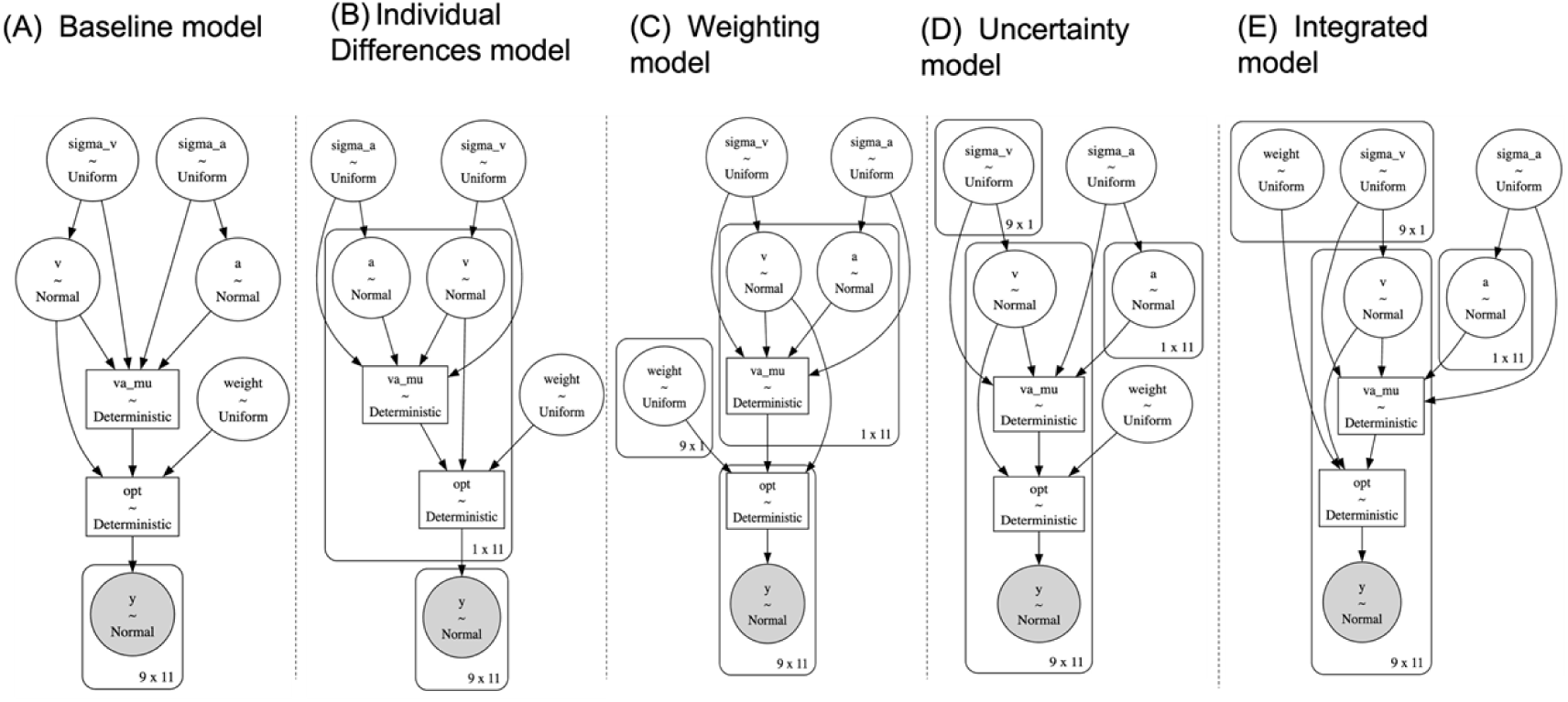
Diagram of Five Bayesian Model Structures. Key elements of the structure diagram: Nodes: Each ellipse represents a random variable or deterministic variable. Nodes filled in gray typically represent observed data. Arrows: Arrows indicate dependency relationships from one variable to another. Deterministic Nodes: Nodes labeled “Deterministic” indicate that the variable is a deterministic function of its parent variables, with values determined by the parent nodes. Shape: Numbers next to nodes represent the shape of tensor variables. For example, 9×1 may represent a vector containing 9 elements. Distribution: For each node that is sampled from a distribution, the specific distribution is provided. For example, “Normal” indicates a normal distribution, “Uniform” indicates a uniform distribution. The parameters of a distribution may depend on other nodes.

Model 1 serves as the baseline model, assuming neither individual differences in unisensory information nor differences across eccentricities. Consequently, each parameter yields only a single optimized value, with all factors considered constant. Model 2 is an individual differences model that, compared to the baseline, accounts for inter-subject variability in sampled sensory information by introducing shape parameters. This allows the model to independently sample v and a for each subject, modeling based on their own sensory information, but still without considering eccentricity effects. Model 3 is a weight model that introduces shape parameters for weight w, enabling separate sampling and estimation of integration weights for each eccentricity, resulting in different estimates across eccentricities. Model 4 is an uncertainty model that assigns shape parameters to *σ*_v_, positing that subjects exhibit different perceptual uncertainties for visual stimuli presented at different spatial locations, leading to eccentricity-dependent differences in subsequent integration. Model 5 is the integrated model, which combines features of the two previous eccentricity-based models. This most complex model aims to simultaneously capture the effects of eccentricity on both weight w and visual information uncertainty *σ*_v_.

Each of the five models was sampled using four parallel MCMC chains, with each chain drawing 2000 samples and discarding the first 1000 for tuning. For the integrated model, which has more parameters and thus requires more samples, each chain drew 4000 samples while discarding the first 2000. The sampling results showed adequate convergence for all parameters across models, with r̄ ≤ 1.01,indicating that the data structure has been thoroughly explored and the models have been adequately fitted to the existing data. *r̄* represents the potential scale reduction factor, which assesses MCMC sampling quality by examining the ratio of between-chain variance to within-chain variance to evaluate whether chains have converged to the same distribution; values close to 1 indicate good convergence, and all models in this experiment achieved stable convergence.

To compare the performance of the five models, we employed Widely Applicable Information Criterion (WAIC) as the evaluation metric. This is a widely used model selection criterion in Bayesian statistics that identifies models fitting the data well without excessive complexity. Its calculation is based on the log pointwise predictive density (*lppd*) and the effective number of parameters (Wasserman, 2000). The specific formula is as follows:

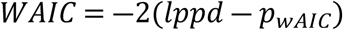

where *lppd* is the expected value of the log-likelihood function of the observed data over the posterior distribution of model parameters, representing goodness-of-fit; and *p*_*wAIC*_ represents the effective number of parameters, estimating model complexity. These two components separately assess model fit and complexity. WAIC seeks to balance these aspects to select the optimal model. The smallest WAIC value typically corresponds to the best model (Table 4).

**Table 4.** Performance Metrics for Different Models.

| Models | $lppd$ | $SE(lppd)$ | $p_{wAIC}$ | $WAIC$ | $\Delta WAIC$ |
| --- | --- | --- | --- | --- | --- |
| Baseline | -30.36 | 6.17 | 1.54 | 63.80 | 65.94 |
| Individual differences | -19.97 | 6.55 | 11.00 | 61.94 | 64.08 |
| Weight | 10.83 | 3.69 | 9.76 | -2.14 | \ |
| Uncertainty | 4.01 | 3.43 | 14.07 | 20.12 | 22.26 |
| Integrated | 7.70 | 3.38 | 13.26 | 11.12 | 13.26 |

We calculated WAIC for the five models and plotted the comparison, shown in Figure 10. The weight model performed best, substantially outperforming all other models, even surpassing the integrated model that considered both uncertainty and weight variations. Therefore, we can conclude that the eccentricity effect obtained in the current experiment operates by directly altering the information weight for integrated audiovisual same-source stimuli, making visual estimates in multisensory contexts more susceptible to interference from auditory information.

**Figure 10.**
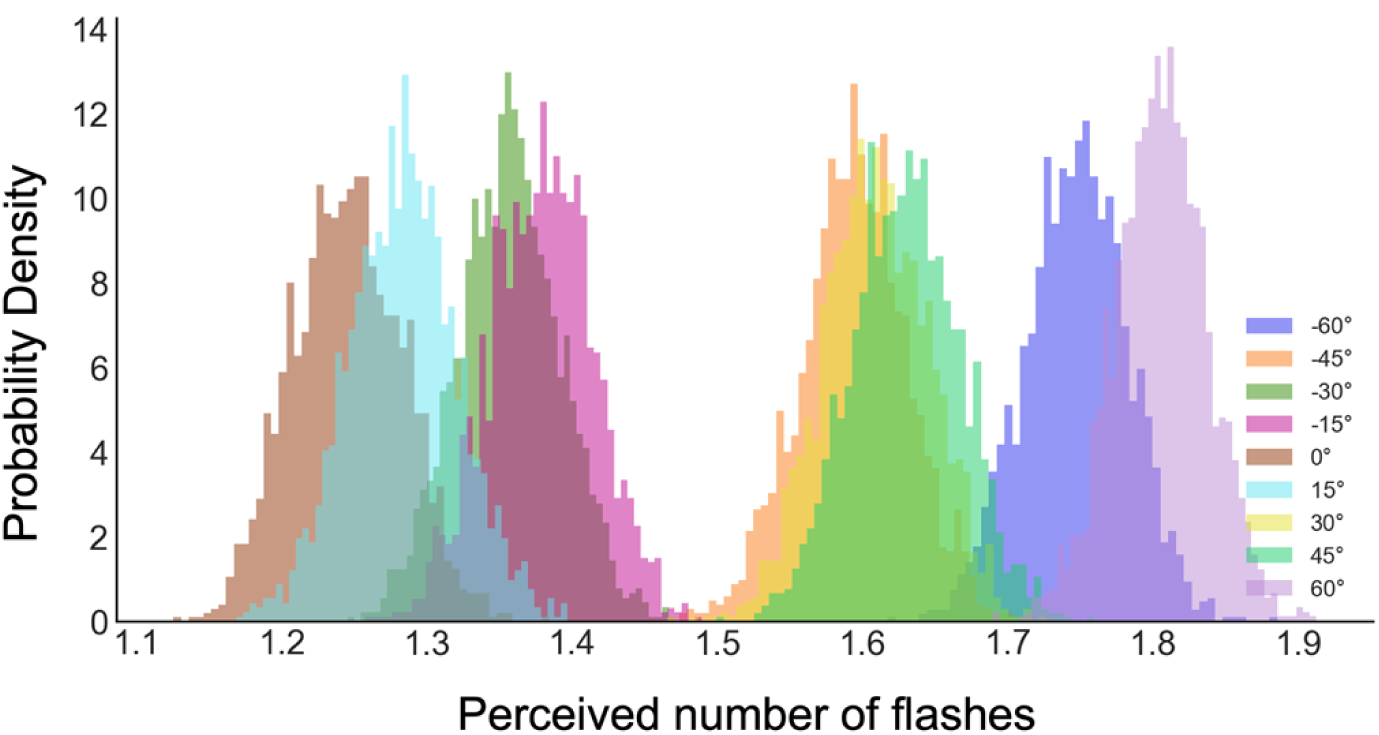
Model-predicted probability density distributions of reported counts. Colors indicate the posterior distributions for different spatial eccentricities. While the means shift across positions, the standard deviations of the distributions remain consistent across conditions.

Table 5 shows the fit of the weight model on specific parameters. The key parameters in this model are *σ*_v_, *σ*_a_, and the respective weights w for each of the 9 different eccentricities. The highest density interval (HDI) is used to represent the posterior distribution, encompassing the region of highest posterior density; here, the 3% and 97% percentiles are used as the lower and upper bounds. *r̄* being close to 1 indicates that the chains have converged well; here, the model’s estimates for each parameter are visible, demonstrating that model sampling has achieved stable convergence based on the existing data.

**Table 5:** Posterior Parameter Distributions of the Weight Model.

| Parameters | $M$ | $SD$ | 3% HDI | 97% HDI | $\bar{r}$ |
| --- | --- | --- | --- | --- | --- |
| $\sigma_v$ | 0.093 | 0.052 | 0.010 | 0.181 | 1.0 |
| $\sigma_a$ | 0.304 | 0.096 | 0.149 | 0.487 | 1.0 |
| $w(-60^\circ)$ | 0.830 | 0.096 | 0.672 | 1.000 | 1.0 |
| $w(-45^\circ)$ | 0.659 | 0.103 | 0.465 | 0.852 | 1.0 |
| $w(-30^\circ)$ | 0.391 | 0.091 | 0.223 | 0.566 | 1.0 |
| $w(-15^\circ)$ | 0.416 | 0.092 | 0.246 | 0.590 | 1.0 |
| $w(0^\circ)$ | 0.265 | 0.090 | 0.097 | 0.436 | 1.0 |
| $w(15^\circ)$ | 0.308 | 0.091 | 0.140 | 0.484 | 1.0 |
| $w(30^\circ)$ | 0.675 | 0.101 | 0.484 | 0.868 | 1.0 |
| $w(45^\circ)$ | 0.699 | 0.102 | 0.511 | 0.892 | 1.0 |
| $w(60^\circ)$ | 0.894 | 0.078 | 0.756 | 1.000 | 1.0 |

To further evaluate the model’s predictive performance, the probability density distribution of the optimal estimate (opt) obtained at each eccentricity level is plotted in Figure 10. It can be observed that as eccentricity extends from central vision toward the periphery, the observer’s estimate of the number of flashes also increases.

Next, we quantitatively describe the mapping between the weighting model’s predicted results (opt) and spatial eccentricity. Assuming that incorrect responses in the 1F2B (1-flash, 2-beep) task consist of reporting two flashes, let *p* denote the response accuracy. Thus, the mean reported number of flashes is N̄ = p +2 × (1 ― *p*) = 2 ― *p*, or conversely, p̄ = 2 ― N. In the behavioral data, we found that a Gaussian distribution effectively fits the spatial distribution of response accuracy. To validate the weighting model’s descriptive power, we calculated a hypothetical accuracy (2 ― opt) and fitted a Gaussian curve to examine whether the model-generated posterior data exhibit spatial characteristics similar to the empirical accuracy distribution.

As illustrated in Figure 11, the optimal estimates predicted by the model also follow a Gaussian relationship across space, R^2^= 0.863, closely mirroring the overall distribution of the actual data. Due to occasional trials where participants reported three flashes (N>2), the actual *p* is slightly lower than the theoretical 2−N. Consequently, the calculated p̄ distribution is slightly higher than the actual response accuracy; nonetheless, the high degree of similarity between the two distributions confirms that the weighting model provides an optimal representation of the experimental data.

**Figure 11.**
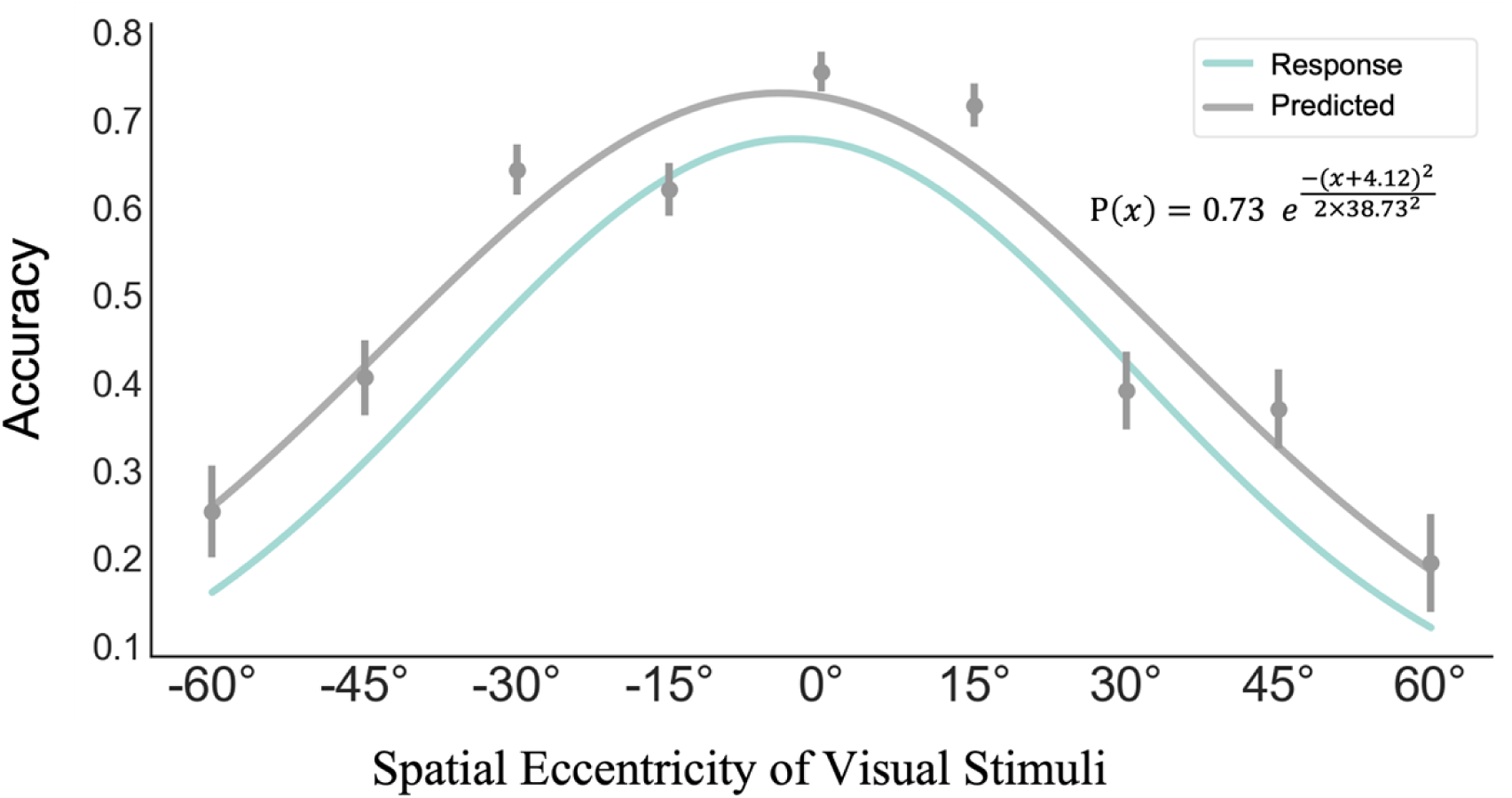
Comparison of model-predicted and empirical accuracy across spatial eccentricities. Gray data points and lines represent the predicted accuracy p̄ calculated from model-sampled opt. The green line (consistent with Figure 8) shows the Gaussian fit of the participants’ empirical response data. Error bars denote ±1 standard error of the mean (SEM).

#### Discussion

Extending Experiment 1, the present experiment documented a robust eccentricity-dependent SIFI up to 60°: the farther into the periphery, the more flashes participants reported in 1F2B trials. Crucially, this spatial modulation emerged only when audition and vision conflicted; unimodal vision and congruent AV trials were flat across eccentricity. Curve-fitting showed that hit-rate follows a Gaussian profile centred on the fovea. Bayesian model comparison revealed that a *weight* model—where the prior probability of fusing AV signals increases with retinal eccentricity—outperformed an *uncertainty* model and a full model that varied both weight and visual noise. Thus, stimuli located more peripherally are a priori more likely to be bound with concurrent sounds, supporting recent proposals that early sensory cortices exhibit space-specific cross-modal weighting (Eckert et al., 2008; Falchier et al., 2002; Rockland & Ojima, 2003).

Predictions generated by the weight model reproduced the empirical Gaussian spatial signature (R² = .86), confirming its explanatory power. In traditional causal-inference accounts, the fusion prior is usually treated as constant because stimulus location is fixed (Shams et al., 2005). Here, freeing the weight parameter captured the spatial prior: observers expect peripheral visual events to be auditory-causal, so auditory input dominates the final estimate. The failure of the uncertainty model aligns with the behavioural null-effect in unisensory flash trials: when attention is equated across locations, visual numerosity perception is spatially uniform, indicating that the visual system can compensate for lower peripheral acuity under unisensory conditions (Shulman et al., 1985).

Our findings converge with M/EEG studies identifying early-latency signatures of the flash illusion (47–120 ms) (Mishra et al., 2008; Shams et al., 2005) and fMRI evidence showing heightened recruitment of the superior temporal sulcus (STS) and superior colliculus (SC) during illusory trials (de Haas et al., 2012). Collectively, these data support the view that multisensory integration is not limited to high-level association cortices; rather, it is an early-stage process automatically modulated by the spatial receptive-field architecture of primary sensory areas.

### 2.3 Experiment 3 – Impact of Spatial AV Congruency on SIFI

After establishing that visual eccentricity biases AV integration, we asked whether auditory spatial position matters. We simultaneously manipulated the location of flashes and beeps to compare integration when AV signals were spatially congruent versus incongruent. Because vision is the dominant modality in SIFI (Kumpik et al., 2014), we expected a robust illusion under both arrangements with no additional penalty for spatial mismatch.

#### Participants

Ten undergraduates (6 female, 60 %; age 19–22, M = 20.50, SD = 0.97) with normal hearing and normal/corrected vision participated. All were right-handed and naïve to the purpose. They received 50 RMB after a 40-min session.

#### Apparatus and Stimuli

Testing took place in the same dimly lit laboratory. Visual stimuli were presented on a 27-in LCD (1920 × 1080, 120 Hz, 56 cm width) viewed at 60 cm. A white Gaussian-ramped disk (1° dia.) appeared 7° left or right of fixation; a 2° white square frame served as location cue. Auditory stimuli were 10-ms, 2000-Hz pure tones delivered via closed-back headphones at 44.1 kHz. Responses were made with “Z” and “?/” keys.

#### Design and Procedure

This experiment aimed to investigate the impact of audiovisual spatial congruency on multisensory integration. Visual stimuli were presented at one of two possible spatial locations (7° eccentricity in the visual field), consisting of one, two, or three flashes. The three-flash condition was included to prevent participants from adopting specific response strategies or developing a distinct response bias in their number judgments.

Catch trials (attention checks) were implemented with no flash present; participants who committed more than five errors in these trials were to be excluded. Notably, all participants in this study committed four or fewer errors. Auditory stimuli consisted of zero to four beeps. Unlike previous experiments that utilized only binaural presentation, here we incorporated spatial cues for the auditory stimuli: beeps were presented either binaurally or monaurally (ipsilateral or contralateral to the visual stimulus).

The trial distribution is detailed in Table 5. As with previous experiments, a higher number of trials were allocated to the critical 1F2B (1-flash, 2-beep) and 2F1B (2-flash, 1-beep) conditions to manipulate the interstimulus interval (ISI) and explore the temporal dynamics of integration. The ISI was set at 42 ms for flashes and 30 ms for beeps (the latter corresponding to the 40 ms Stimulus Onset Asynchrony (SOA) used in Experiments 1 and 2). For the 1F2B and 2F1B conditions, four distinct temporal relationships were designed based on the stimulus sequence and interval length. Specifically, the “long interval” was three times the duration of the “short interval”. This variety in temporal combinations was designed to increase experimental diversity and facilitate a preliminary investigation into the combined effects of spatial congruency and temporal context on integration. Consequently, the number of 1F2B and 2F1B trials was four times that of other conditions. The experiment comprised 684 trials in total, with mandatory breaks of at least one minute every 50 trials. The total duration was approximately 40 minutes.

**Table 5.** Trial distribution across conditions for Experiment 3.

|  |  | Number of auditory stimuli |  |  |  |  |  |
| --- | --- | --- | --- | --- | --- | --- | --- |
|  |  | 0 | 1 | 2 | 3 | 4 | Sum |
| Number<br>of<br>flashes | 0 | 6 | 18 | 18 | 18 | 18 | 78 |
|  | 1 | 12 | 36 | 144 | 36 | 36 | 264 |
|  | 2 | 12 | 144 | 36 | 36 | 36 | 264 |
|  | 3 | 6 | 18 | 18 | 18 | 18 | 78 |

The procedure was largely consistent with those of Experiments 1 and 2. Each trial began with a white fixation cross presented at the center of a gray screen for 500 ms. Subsequently, a white square (2° in visual angle) appeared for 500 ms on either the left or right side to cue the spatial location of the upcoming target. After a 500 ms blank-screen interval, the audiovisual stimuli were presented.

Following the stimulus presentation, the fixation cross turned red, serving as a “go” signal for the participant to respond. Once a response was made, the screen transitioned to a blank display. Participants reported seeing one or two flashes by pressing the “Z” or “/” keys, which were counterbalanced across participants. In cases where three flashes were perceived, participants were instructed to press the spacebar. The maximum response window was 2.5 s. The inter-trial interval (ITI) was 0.75 s, with an additional random jitter incorporated to sufficiently sample various response states of the participants.

#### Data analysis

Similar to Experiment 2, all response data from this experiment were aggregated, and the mean reported numbers under various conditions were calculated. Comparisons across levels were conducted using repeated measures ANOVA, with additional Bayesian factor analysis performed in JASP (JASP Team, 2023).

First, we examined whether a significant SIFI effect was observed by calculating the mean reported numbers for all participants under each auditory stimulus condition when visual stimuli were 1 and 2 flashes, with results shown in Figure 12. A two-way repeated measures ANOVA was conducted. Since the auditory stimulus number level failed Mauchly’s test of sphericity, χ²(9) = 29.870, p < .001, Greenhouse-Geisser correction was applied. The data revealed a significant main effect of flash number on participants’ reports, with reports under two flashes significantly higher than under one flash, F(1, 9) = 76.192, p < .001, ηp² = .894, BF₁₀ = 2615.170, mean difference |MD| = 0.544. The main effect of auditory stimulus number was also significant, F(1.4, 12.597) = 28.908, p < .001, ηp² = .894, BF₁₀ = 1.437 × 10⁸. The interaction between the two factors was significant as well, F(4, 36) = 2.647, p = .049, ηp² = .227, BF₁₀ = 1.419.

**Figure 12.**
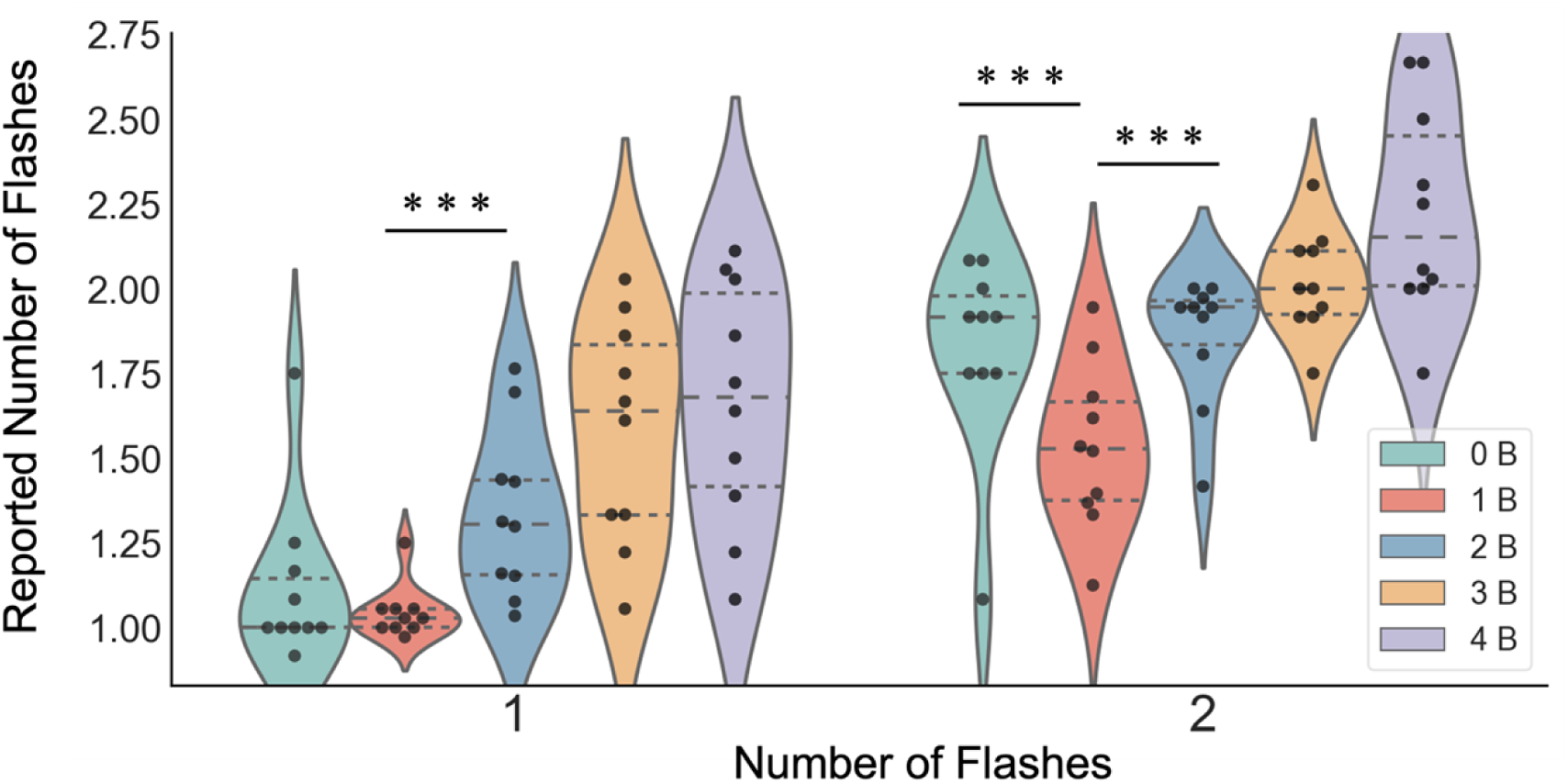
Violin plot of participants’ reported numbers in Experiment 3. The width of the violin plot represents the density distribution of the data; each point represents the data of one participant under that condition. The horizontal lines inside the violin plot represent the upper quartile, median, and lower quartile of the data. * indicates p < .05, ** indicates p < .01, *** indicates p < .001.

Focusing specifically on the simple main effects of auditory stimuli under the two flash levels: When the target flash was 1 flash, the simple main effect of auditory number was significant, F(4, 36) = 19.461, p < .001, BF₁₀ = 1.560 × 10⁶. Specifically, no significant difference existed between 1B and 0B conditions, |MD| = 0.072, p = 1.000, BF₁₀ = 0.586. However, with more than 1 auditory stimulus, participants’ mean reported numbers were significantly higher than both the no-auditory-stimulus and audiovisual-congruent conditions, ps < .001, BF₁₀ > 1.5. When the target flash was 2 flashes, the simple main effect of auditory number was also significant, F(4, 36) = 23.301, p < .001, BF₁₀ = 1.108 × 10⁷. Participants’ reported numbers in the 1B condition were significantly lower than in other conditions, ps < .010, BF₁₀ > 25.

That is, this experiment observed both significant flash fission illusion and flash fusion illusion: When auditory stimuli presented more stimuli than visual flashes, participants’ reported numbers increased significantly, and when presented auditory stimuli were fewer than target flashes, participants’ reported numbers were significantly lower than other conditions and unimodal perception without auditory interference. Notably, although the three experiments so far have only found illusory interference during audiovisual incongruence without observing significant facilitation during consistent audiovisual information, in this experiment, the variance across all participants’ data was notably smaller under the 1F1B condition, suggesting that congruent conditions may facilitate more efficient processing of visual information.

To further examine the influence of spatial features of audiovisual stimuli on integrated perception, both fission and fusion illusions were observed in this experiment. However, the fission illusion phenomenon was more pronounced, with multiple conditions triggering fission perception. Therefore, the mean of participants’ reports when the target flash was presented once was adopted as the comparison metric.

To avoid potential influences from visual presentation field, a two-way ANOVA of spatial location and auditory stimulus number was first conducted. As shown in the left panel of Figure 13, the main effect of spatial field on participants’ reported numbers was not significant; presenting stimuli in the left versus right visual fields had no effect on audiovisual integration, F(1, 9) = 0.056, p = .819, ηp² = .006, BF₁₀ = 0.347. Therefore, when subsequently comparing audiovisual stimulus congruence, trials presented in the left and right visual fields were combined, focusing only on the relative spatial relationship between audiovisual stimuli.

**Figure 13.**
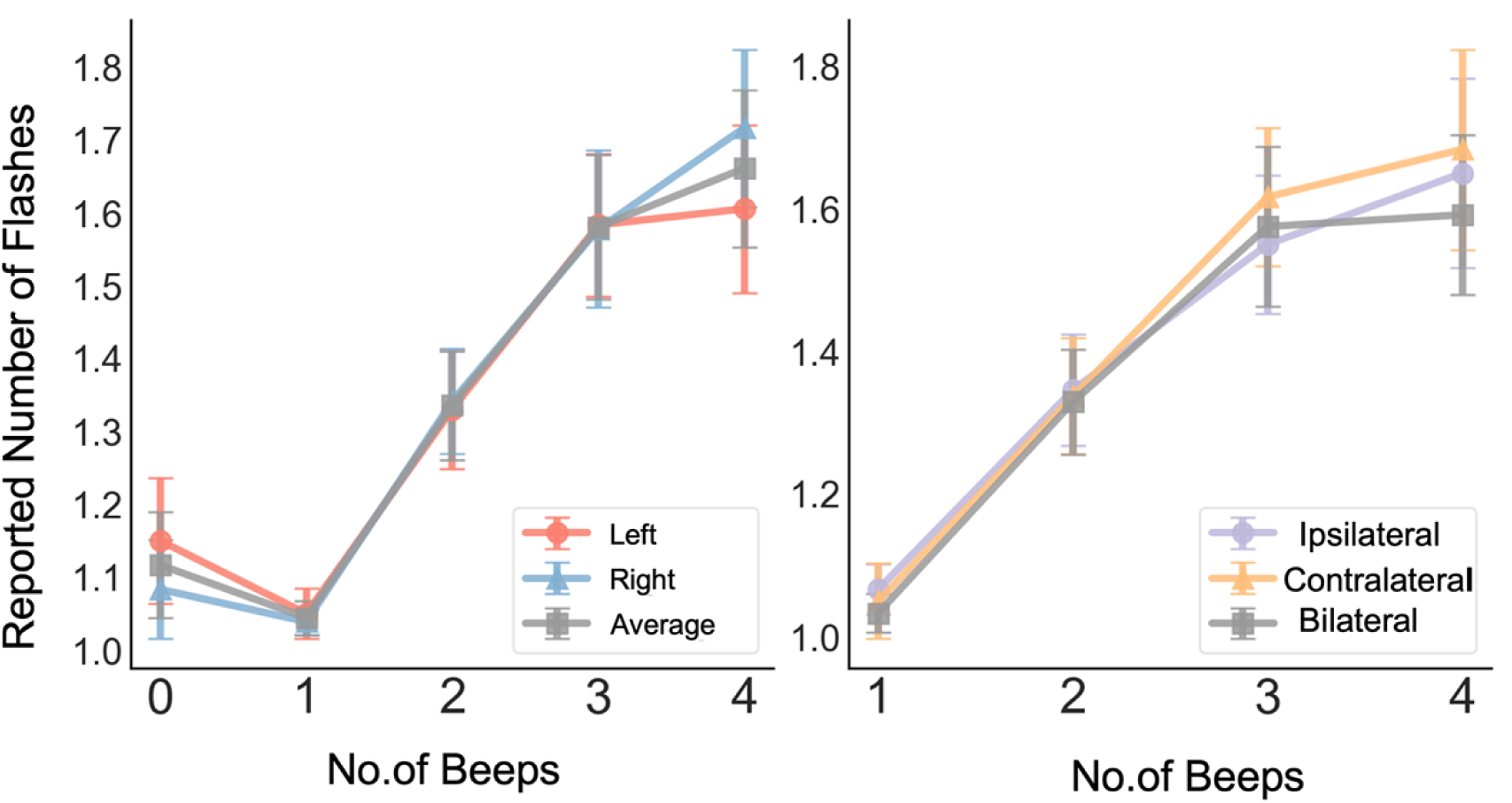
Participants’ reported numbers under different conditions in Experiment 3. The left panel shows the reported number of flashes under different auditory stimulus numbers when visual stimuli were presented in the left or right visual field. The right panel combines stimuli from both visual fields, comparing reported numbers when auditory stimuli were presented ipsilaterally, contralaterally, and bilaterally to the visual target. * indicates p < .05, ** indicates p < .01, *** indicates p < .001.

Participants’ reported numbers were then compared when auditory stimuli were presented ipsilaterally, contralaterally, or bilaterally to the flash. ANOVA results showed that audiovisual stimulus congruence had no significant effect on participants’ perceived numbers. Whether the sound was presented ipsilaterally, contralaterally, or bilaterally to the visual stimulus, participants reported similar numbers of flashes, F(2, 18) = 0.494, p = .618, ηp² = .052, BF₁₀ = 0.170.

Thus, it can be concluded that in audiovisual integration paradigms based on visual tasks such as the sound-induced flash illusion, the spatial location of auditory stimuli and the spatial relationship between auditory and visual stimuli have minimal influence on observers’ perception.

#### Discussion

This experiment focused on the spatial congruence of audiovisual stimuli and revealed that the perception of SIFI was not significantly influenced by the spatial relationship between the auditory and visual stimuli. Specifically, whether the visual stimulus was presented in the left or right visual field, and whether the auditory stimulus was presented ipsilaterally, contralaterally, or neutrally (binaurally) to the visual stimulus, no significant effect on participants’ SIFI perception was observed. This finding suggests that spatial congruence of stimuli does not have a measurable effect on the level of audiovisual integration with SIFI.

Notably, in addition to exhibiting the fission illusion—similar to the first two experiments—this experiment also demonstrated a pronounced fusion illusion when the number of auditory stimuli was fewer than the number of flashes. The main distinction between this experiment and Experiment 1 lies in the inclusion of additional stimulus conditions at the audiovisual level to mitigate potential systematic errors. When an individual’s prior perception regarding the overall possible number of audiovisual stimuli in the experiment changes, the level of audiovisual integration is subject to the influence of this perceptual expectation (Wang et al., 2019). This shift in expectation may account for the observed differences in participant performance, even under similar temporal and spatial conditions.

In contrast to the findings of the first two experiments, the results of the current experiment provide greater support for a late-integration model of audiovisual processing. The observation that the SIFI effect under spatially incongruent (contralateral) conditions was similar to the effect under binaural presentation suggests that audiovisual integration may be mediated by higher-level brain regions involved in later-stage processing.

## 3 General Discussion

This study exploited the robustness of the Sound-Induced Flash Illusion (SIFI) to investigate the effect of stimulus spatial characteristics on audiovisual integration capacity, successfully revealing the distribution pattern of how the level of audiovisual integration is influenced by stimulus spatial features, while replicating the SIFI originally discovered by Shams et al. (2002)(Shams et al., 2002).

In Experiment 1 and Experiment 2, we explored the influence of visual stimulus spatial location on observers’ perception during audiovisual integration. The results indicated that, after controlling for matched attentional resource allocation across spatial locations, unimodal visual perception did not change with the spatial location of the stimulus. However, a clear difference in audiovisual bimodal processing was found between spatial locations: stimuli presented in the peripheral visual field (beyond 15°) were more significantly interfered with by auditory information, and the susceptibility to audiovisual integration increased towards the periphery, a spatial distribution that can be approximated by a Gaussian curve. With modeling, we found that the unimodal information uncertainty in the peripheral visual field did not, in fact, change. Instead, the effect was driven by a mechanism that directly enhanced the influence of auditory information by altering the weighting ratio of audiovisual information during processing. In Experiment 3, we attempted to simultaneously manipulate the spatial location of both visual and auditory stimuli to explore whether their spatial congruence would affect the probability of integration. The results showed that SIFI remained stable across various spatial relationships between audiovisual stimuli; even when a potential interhemispheric integration (with contralateral layout of audiovisual stimuli) of sensory information was required, participants’ perceptual performance was virtually identical to conditions with ipsilateral or binaural auditory stimulus presentation. With three experiments, we comprehensively explored the spatial characteristics of audiovisual integration from two aspects: the spatial location of visual information and the spatial relationship between audiovisual stimuli.

This work addresses existing gaps in the literature by expanding the range of spatial eccentricity to a broad 60° across both hemifields while strictly controlling for attentional consistency. These refinements provide a more robust resolution to previously disputed questions regarding spatial modulation of the SIFI. Furthermore, by integrating Bayesian modeling with Causal Inference theory, we characterize the underlying mechanisms from a computational perspective. Our analysis suggests that different retinotopic locations possess intrinsic differences in their responsiveness to audiovisual stimuli, which directly modulates the informational weighting during perceptual inference. This indicates that cross-modal influence is shaped by stimulus location in a relatively automatic, bottom-up manner (Keil & Senkowski, 2018).

There remain certain limitations in the current experiments. For instance, the imbalance in trial design—the deliberate inclusion of more audiovisual conflict trials (1F2B or 2F1B)—could raise concerns about the validity of the conclusions. Such an overall design might lead participants to form a certain expectation regarding the corresponding combinations, e.g., higher accuracy for perceiving two flashes when one auditory stimulus is presented. Previous studies have shown that such perceptual expectation can influence the probability of SIFI (Wang et al., 2019).

To minimize the impact of trial imbalance, we informed participants that all stimulus combinations were possible and utilized Bayes Factor analysis (Dienes, 2014) to robustly compare hypotheses across unequal sample sizes. Because the trial distribution was uniform across all eccentricities, our primary spatial comparisons remain valid. Furthermore, the robust illusions observed contradict the notion that trial frequency awareness suppressed cross-modal influence (Wang et al., 2019). The design strategically maximized sampling of illusory trials while variable SOAs prevented practice effects. Future research should employ neuroimaging to localize these effects in unimodal sensory cortices and incorporate eye-tracking to control for microsaccades and bottom-up attentional capture.

In summary, this study provides a comprehensive characterization of the spatial constraints governing audiovisual integration. Behaviorally, we refined previous explorations by demonstrating that SIFI susceptibility is significantly modulated across a broad 60°range of visual eccentricity. Conversely, our investigation into spatial incongruence revealed that the relative position of auditory stimuli does not significantly influence integration, highlighting the dominance of visual eccentricity in shaping these percepts. At the modeling level, a Gaussian distribution successfully quantified perceptual performance across the visual field, providing a robust mathematical description of these spatial variations. Furthermore, Bayesian computational modeling localized the eccentricity effect to a fundamental shift in the allocation of sensory weights: visual stimuli in the far periphery possess a higher probability of being integrated with auditory information compared to those at fixation. By systematically manipulating spatial characteristics within the flash illusion paradigm, this research deepens our understanding of the mechanisms underlying multisensory processing. These findings offer a critical empirical framework for the selection of stimulus locations in future multisensory research and contribute to a more nuanced model of how the brain resolves audiovisual information across the visual field.

